# TGFβ1-TNFα regulated secretion of neutrophil chemokines is independent of epithelial-mesenchymal transitions in breast tumor cells

**DOI:** 10.1101/2024.10.11.617845

**Authors:** Shuvasree SenGupta, Erez Cohen, Joseph Serrenho, Kaleb Ott, Pierre A. Coulombe, Carole A. Parent

**Affiliations:** Life Sciences Institute, University of Michigan, Ann Arbor, MI; Department of Pharmacology, University of Michigan Medical School, Ann Arbor, MI; Department of Cell & Developmental Biology, University of Michigan Medical School, Ann Arbor, MI; Undergraduate Research Opportunity Program, University of Michigan, Ann Arbor, MI; Rogel Cancer Center, University of Michigan Medical School, Ann Arbor, MI; Department of Dermatology, University of Michigan Medical School, Ann Arbor, MI

## Abstract

Neutrophils have tumor-promoting roles in breast cancer and are detected in higher numbers in aggressive breast tumors. How aggressive breast tumors recruit neutrophils remains undefined. Here, we investigated the roles of TGF-β1 and TNF-α in the regulation of neutrophil recruitment by breast cancer cells. TGF-β1 and TNF-α are pro-inflammatory factors upregulated in breast tumors and induce epithelial to mesenchymal transitions (EMT), a process linked to cancer cell aggressiveness. We report that, as expected, dual treatment with TGF-β1 and TNF-α induces EMT signatures in premalignant M2 cells, which are part of the MCF10A breast cancer progression model. Conditioned media (CM) harvested from M2 cells treated with TGF-β1/TNF-α gives rise to amplified neutrophil chemotaxis compared to CM from control M2 cells. This response correlates with higher levels of the neutrophil chemokines CXCL1, CXCL2, and CXCL8 and is significantly attenuated in the presence of a CXCL8-neutralizing antibody. Furthermore, we found that secretion of CXCL1 and CXCL8 from treated M2 cells depends on p38MAPK activity. By combining gene editing, immunological and biochemical approaches, we show that the regulation of neutrophil recruitment and EMT signatures are not mechanistically linked in treated M2 cells. Finally, analysis of publicly available cancer cell line transcriptomic databases revealed a significant correlation between CXCL8 and TGF-β1/TNF-α-regulated or effector genes in breast cancer. Together, our findings establish a novel role for the TGF-β1/TNF-α/p38 MAPK signaling axis in regulating neutrophil recruitment in breast cancer, independent of TGF-β1/TNF-α regulated EMT.

## INTRODUCTION

The early and prominent role of neutrophils in response to infections or injuries is well-established. In addition, the broad-spectrum functionality of neutrophils as part of the tumor niche, with a significant impact on tumor progression and metastasis, has been emerging for a broad array of malignancies (1, 2). In breast cancer, for instance, neutrophils and neutrophil-derived products are more frequently detected in the tumor niche of the highly aggressive triple-negative breast cancer subtype (TNBC) compared to the less aggressive hormone receptor-positive (HR^+^) subtype (3–5) and have been shown to promote cancer cell metastasis (3, 4). When comparing the neutrophil recruiting abilities of cancer cell lines representing different breast cancer subtypes, we showed that tumor-conditioned media (TCM) isolated from TNBCs induce greater neutrophil chemotaxis relative to TCM from HR^+^ breast cancer cells. We also found that CXCR2 ligands and transforming growth factor-β1 (TGF-β1) secreted from TNBC cells work in tandem to recruit neutrophils (6). These findings indicate a link between cancer cell aggressiveness and neutrophil recruiting capabilities.

A well-established response linked to cancer cell aggressiveness is the epithelial-to-mesenchymal transition (EMT) (7–9). EMT is a complex cellular process where epithelial cells within a tissue lose polarity and cell-cell adhesion and gain invasive properties as they adopt a mesenchymal character. This transition is dynamic, with cells oscillating between interconvertible epithelial and mesenchymal states and expressing varying degrees of EMT signature markers. EMT promotes cell motility, which plays a crucial role in mediating aggressiveness and metastasis (10–13). In addition, positive correlations between EMT signatures and the composition of tumor-infiltrated immune cells have been observed in various tumors (14, 15). However, whether EMT is directly responsible for regulating immune cell recruitment to tumors remains an open issue. It has been established that EMT signatures are induced when epithelial cells are exposed to inflammatory mediators commonly encountered in tumor niches, such as cytokines, chemokines, and growth factors (7, 16). The EMT-inducing effects of such inflammatory mediators are regulated by the activation of signaling pathways that induce the expression of classical EMT transcription factors (TFs) including snail, twist, and zeb (17–20). These EMT-driving TFs then control the expression of several mesenchymal signature markers such as vimentin (Vim), N-cadherin (N-Cad), and fibronectin (Fn), and epithelial signature marker E-cadherin (E-Cad) (21, 22). In addition, activation of signaling pathways, for instance SMAD and nuclear factor κ-light-chain enhancer of activated B cells (NF-κB), can directly regulate the expression of EMT signature markers independently of the classical EMT TFs (23–26).

TGF-β, which belongs to a multi-functional cytokine family, is a well-established EMT inducer (21). TGF-β exists in three isoforms, TGF-β1, TGF-β2, and TGF-β3, with TGF-β1 being the most prevalent in many cancer types (27). TGF-β1 signals through the TβRI/TβRII heterotetrameric receptor complex, resulting in the activation of canonical SMAD-dependent and multiple non-canonical, SMAD-independent signaling pathways that include c-Jun NH2-terminal kinase (JNK), p38 mitogen activated protein kinase (MAPK), extracellular signal regulated kinase (ERK), phosphatidylinositol 3-kinase (PI3K)-protein kinase B (AKT), and NF-κB (28). TGF-β1 promotes EMT through both SMAD-dependent and -independent pathways (29). Furthermore, TGF-β1 is produced by both cancer cells and stromal cells present in the tumor niche (30), and the presence of TGF-β1 in breast tumors has been associated with increased lymph node metastasis (31). We also found that aggressive TNBC cell lines secrete high levels of TGF-β1 (6). Whether the activation of TGF-β1 signaling pathways impacts the ability of cancer cells to recruit neutrophils to breast tumors remains, however, unknown.

Another EMT inducing factor, tumor necrosis factor-α (TNF-α), is an important pro-inflammatory cytokine that is highly upregulated in breast cancer cells and secreted by stromal cells (32, 33). TNF-α belongs to the TNF/TNFR superfamily of cytokines and binds TNFR1 to activate JNK, MAPK, and NF-κB signaling pathways (34). TNF-α also coordinates with TGF-β to induce EMT in multiple cancer types, including breast cancer (35–38). The combined action of TNF-α and TGF-β1 can potentiate the activation of signaling pathway/s and induce robust EMT signatures and cancer cell invasiveness (36). Yet, how the combined action of TGF-β1 and TNF-α controls the secretion of neutrophil guidance cues in breast tumors has not been studied.

In this study, we used cell lines of the MCF10A breast cancer progression model (39, 40) as well as human breast cancer cell lines (41) to identify the mechanisms by which aggressive breast cancer cells recruit neutrophils. In particular, we assessed the role of TGF-β1/TNF-α treatments and EMT-associated changes on this response and validated our experimental findings by conducting a comprehensive analysis of transcriptomics datasets from breast cancer cell lines. We found that the neutrophil recruitment ability is not regulated by EMT signatures. Instead, these two processes appear to occur in parallel and can be triggered by distinct signaling pathways downstream of TGF-β1/TNF-α.

## RESULTS

### TGF-β1/TNF-α treated M2 cells recruit neutrophils by inducing chemokine secretion

We first used the non-malignant MCF10A cells, the M1 cells in the breast cancer progression model (39, 40) to investigate the effect of TGF-β1/TNF-α treatment on their neutrophil recruitment ability. However, these cells failed to survive when subjected to serum free conditions for the collection of the conditioned media (CM), a step necessary to minimize neutrophil activation by the factors present in the serum (6). We therefore used M2 cells, a H-Ras transformed, pre-malignant derivative of the M1 cells that undergo minimal cell death under serum free condition (39, 40). We first characterized the effect of TGF-β1/TNF-α dual treatment on the morphology of M2 cells. While control, non-treated cells retain a tightly packed, cuboidal and epithelial-like morphology (Fig. 1Ai), treated cells become loosely organized and acquire an elongated, spindle-like shape with front-back polarity (Fig 1Aii), consistent with a mesenchymal phenotype (10, 42). The treated cells also display dramatic changes in actin filament organization and an increase in cytosolic and nuclear areas compared to control cells (Fig.1Aiii,iv; 1Bi,ii). Immunostaining revealed that expression of the mesenchymal markers Vim, N-Cad and Fn are upregulated with TGF-β1/TNF-α dual treatment, consistent with EMT-associated changes (Fig.1Av,vi,vii,viii;1Biv,v,vi). Increases in the total protein levels of N-Cad and Fn were also confirmed by western blotting (Fig. 1Cii,iii). While treated cells retained a similar level of total E-Cad compared to control cells (Fig.1Ci), TGF-β1/TNF-α treatment induced a profound redistribution of E-Cad, from being strictly localized to cell-cell junctions in control cells (Fig.1Av) to a more diffused cytosolic distribution in treated cells (Fig.1Avi), along with a decrease in the cortex to cytosolic ratio of E-Cad intensity (Fig.1Biii).

**Figure 1.**
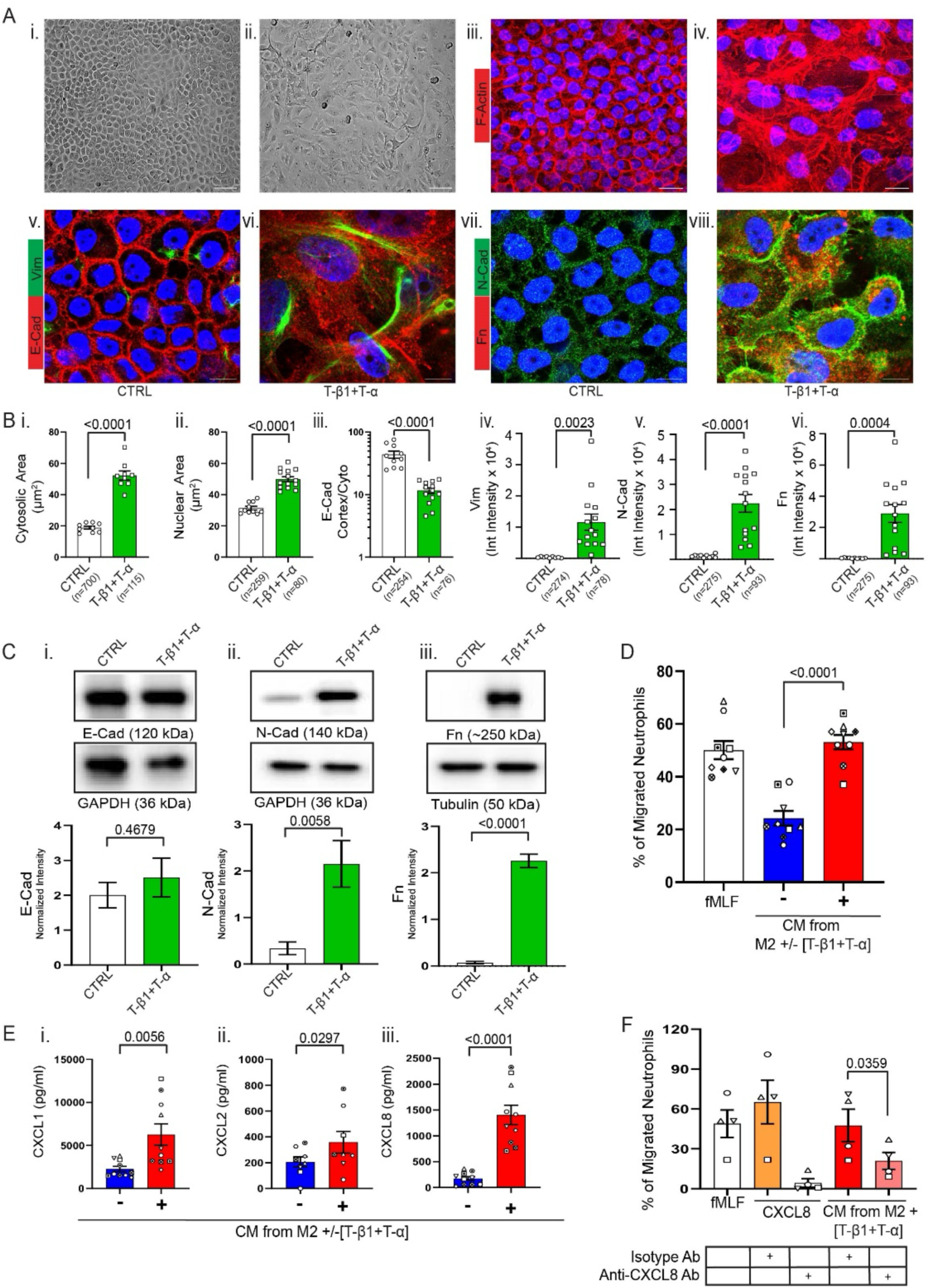
TGF-β1/TNF-α treatment amplifies the neutrophil recruiting activity of M2 cells by inducing chemokine secretion. **A.** Representative bright field (i, ii) and IF images (iii-viii) (n=3) of control (CTRL) M2 cells (i, iii, v, vii) or M2 cells treated with a combination of 20 ng/ml T-β1 (TGF-β1) and 100 ng/ml T-α (TNF-α) [T-β1+ T-α] for 72 hrs (ii, iv, vi, viii). Airyscan confocal microscopy images showing MIP (iii, iv) or a single z image (v-viii) of fixed M2 cells stained for F-actin with phalloidin-TRITC (red) (iii, iv), E-Cad (red)/Vim (green) (v, vi), N-Cad (green)/Fn (red) (vii, viii), and nuclei with DAPI (blue). Scale bar 50 µm for bright filed images and 20 µm (iii, iv) and 10 µm (v, vi, vii, viii) for IF images. **B.** Graphs depicting cytosolic (i) and nuclear (ii) area or cortex/cytosolic intensity ratio of E-Cad (iii) or integrated intensity measures of Vim, N-Cad, Fn (iv,v,vi) in control vs T-β1/T-α treated M2 cells. Each dot represents an average of all cells in each image (≥3 images/condition/experiment). Total number of cells (n) analyzed is reported under each condition. **C.** Top: Representative western blots of the respective markers from CTRL or T-β1/T-α treated cells. Bottom: Graphs showing band intensities of the markers normalized to the loading controls (mean values +/− SEM from n=3). **D.** Graph depicting the percentage of neutrophils that migrated into the bottom chamber of transwells containing equal volume of CM from CTRL or T-β1/T-α treated M2 cells or positive control fMLF (mean values +/− SEM from n=9). Each dot represents the response of neutrophils from an independent donor. **E.** Graphs showing the amount (pg/ml) of CXCL1 (i), CXCL2 (ii), and CXCL8 (iii) secreted by CTRL or T-β1/T-α treated M2 cells (mean values +/− SEM from n=8-10). Each dot represents the value from one experiment. **F.** Graphs showing the percentage of neutrophils that migrated in response to CXCL8 or CM-derived from T-β1/T-α treated M2 and pre-incubated with anti-CXCL8 antibody or isotype control or to control fMLF (mean +/− SEM from n=4). P values were determined using Unpaired t-test (B,C) or repeated measures (RM) One-way analysis of variance (ANOVA) with Dunnett’s multiple comparisons test (D) or Paired t-test (E,F).

We next characterized the neutrophil recruitment ability of CM harvested from control and TGF-β1/TNF-α treated M2 cells. Using a neutrophil transwell migration assay, we found that the CM from treated cells gives rise to a robust neutrophil chemotactic response, comparable to the activity of the formylated bacterial tripeptide fMLF (Fig.1D) - a potent neutrophil chemoattractant (43, 44). To identify the neutrophil chemoattracting factors secreted from treated M2 cells, we first used ELISA and screened for CXCR2 ligands and TGF-β1. While there was undetectable or minimal TGF-β1 present in the CM regardless of treatment (Table S1), we detected >8-fold increase in CXCL8, a 3-fold increase in CXCL1 and a slight increase in CXCL2 levels in the CM from treated cells relative to CM from control cells (Fig.1Ei-iii). Because CXCL8 is a highly potent neutrophil chemokine, we tested whether blocking CXCL8 with a specific neutralizing antibody (Ab) in the transwell system would reverse the effect of CM from treated cells on neutrophil migration. Incubation of CM from treated cells with anti-CXCL8 Ab at a dose that was sufficient to inhibit recombinant CXCL8-induced neutrophil migration significantly reduced the ability of CM from TGF-β1/TNF-α-treated cells to induce neutrophil chemotaxis, compared with the activity of the same CM incubated with a corresponding isotype Ab (Fig.1F). Taken together, these findings establish a role for TGF-β1/TNF-α treatment in inducing changes associated with EMT-like states and in regulating neutrophil chemotaxis by stimulating the release of neutrophil recruiting chemokines, CXCL8 in particular, from M2 cells.

### TGF-β1/TNF-α treated M2 cells recruit neutrophils in a snail- and twist-independent fashion

The EMT-promoting TF snail has been reported to stimulate myeloid cell recruitment to murine tumor models of lung and ovarian cancer (45, 46) and induce neutrophil recruiting chemokine expression in breast cancer cell lines (47). Because the amplified neutrophil recruiting activity of the treated M2 cells is accompanied by EMT-associated changes (Fig.1), we investigated the role of snail and twist, two key EMT-regulating TFs, on neutrophil recruitment. We did so by exogenously expressing an active mutant form of snail (eGFP-snail6SA) tagged with green fluorescent protein (GFP) (48) and twist in M2 cells (Fig.S1A-D) and, in parallel studies, by depleting these TFs from M2 and TNBC cell lines (Fig.S1E,F). While M2 and M2-GFP control cells had low endogenous expression of snail and twist (Fig.S1A,B; see high exposure images), we measured a robust expression of eGFP-snail and twist in M2-eGFP-snail and M2-twist expressing cells (Fig.S1A,B). We also detected a strong nuclear localization of eGFP-snail in M2-eGFP-snail cells and of twist in M2-twist cells (Fig.S1D). Conversely, using shRNA, we obtained a ~70% knockdown of snail in the invasive breast cancer BT549 cells (6, 41) (Fig.S1E) and a depletion of twist in M2, M4, and BT549 twist KO cell lines (Fig.S1Fi-iii).

We found that expression of E-Cad, Fn, and N-Cad is similar in M2-GFP, M2-eGFP-snail, and M2-twist cells in the absence of TGF-β1/TNF-α (Fig.2Ai-iii), suggesting that expression of either snail or twist does not suffice to induce EMT in M2 cells. In agreement with the established role of snail in promoting cell motility (48–51), we also observed a greater proportion of untreated M2-eGFP-snail cells migrating towards serum compared to the M2-GFP or M2-twist cells (Fig.2Bi). Dual treatment with TGF-β1/TNF-α upregulated the expression of Fn and N-Cad, and similarly enhanced the migration of all three cell lines (Fig.2Aii-iii, 2Bii). Surprisingly, we found that TGF-β1/TNF-α treatment of M2-twist KO cells induced a mesenchymal cell-like morphology (Fig.2C) and significantly increased the expression of both Fn and N-Cad (Fig.2D). Together, these findings suggest that snail and twist are dispensable for the expression of EMT markers in M2 cells.

**Figure 2.**
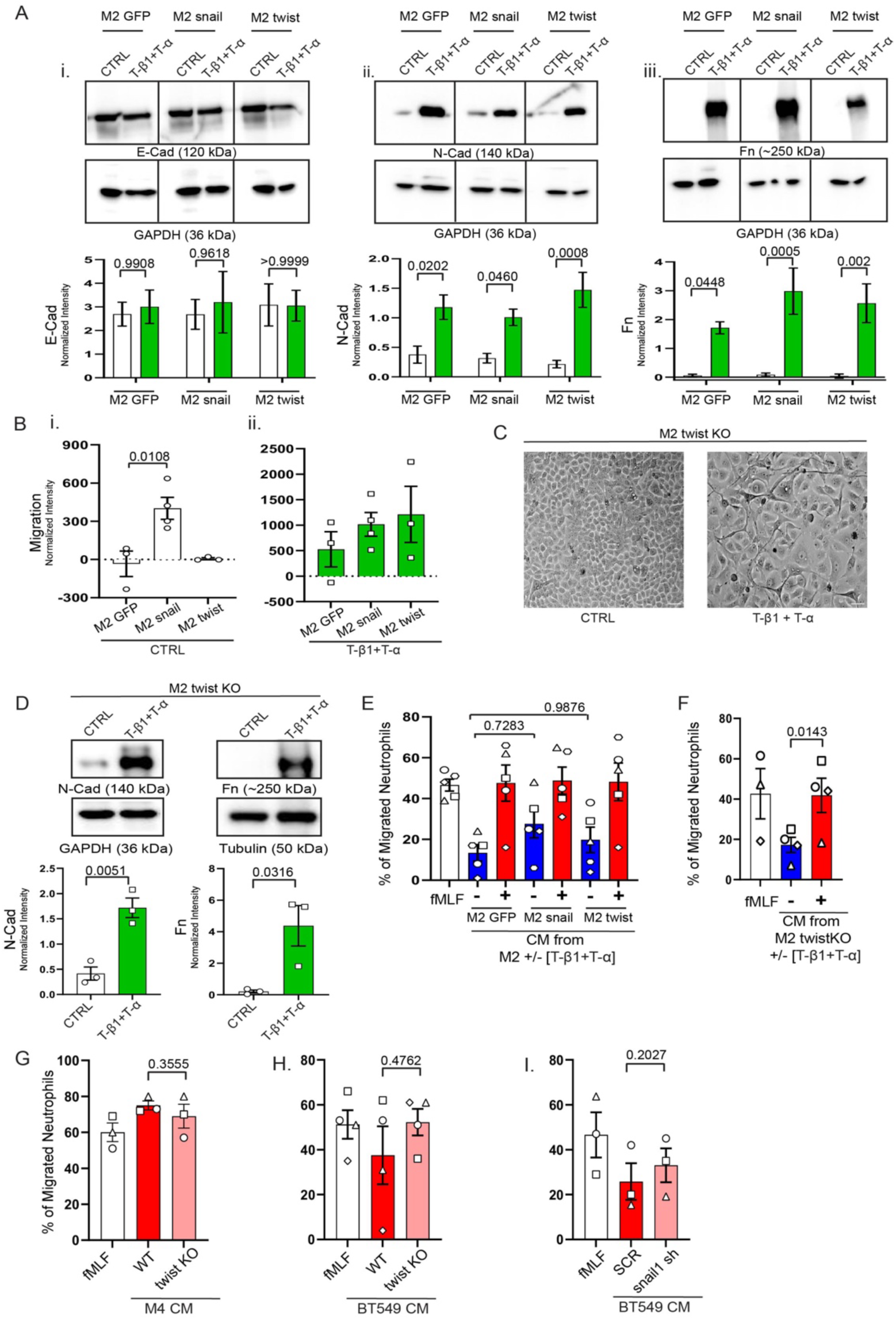
Neutrophil recruitment by TGF-β1/TNF-α treated M2 cells is independent of snail and twist. **A.** Top: Representative western blots showing the expression of (i) E-Cad, (ii) N-Cad, and (iii) Fn in CTRL vs. T-β1/T-α treated cells. Bottom: Graphs showing band intensities of the markers normalized to the respective loading controls (mean values +/− SEM from n=3). **B.** Graphs depicting normalized fluorescence intensity measurements of migrated CTRL or T-β1/T-α treated cells (mean values +/− SEM from n=3-4). **C.** Representative (n=3) bright field images showing morphological changes in M2 twist KO cells with T-β1/T-α treatment. Bar=20 µm. **D.** Top: Representative western blots of N-Cad and Fn expression in M2 twist KO cells. Bottom: Graphs showing band intensities of the markers normalized to the respective loading controls (mean values +/− SEM from n=3). **E-F.** Graphs depicting the percentage of neutrophils that migrated into the bottom chamber of transwells containing equal volume of CM from CTRL or T-β1/T-α treated cells or positive control fMLF (mean values +/− SEM from n=3-5). Each dot represents response of neutrophils from independent donors. **G-I.** Graphs depicting the percentage of neutrophils that migrated into the bottom chamber of transwells containing equal volume of CM from WT or twist KO cell lines for M4 (G) and BT549 (H), and from SCR or snail sh cell line for BT549 (I) or positive control fMLF (mean values +/− SEM from n=3-4). Each dot represents response of neutrophils from independent donors. P values were determined using 2-way ANOVA with Sidak (A) or one-way ANOVA with Dunnett’s (B) or Turkey (E) multiple comparisons test or Unpaired (D) or paired (F-I) t test.

We next evaluated the neutrophil recruiting ability of CM harvested from M2 cells with altered levels of snail and twist expression. We found that CM harvested from M2-eGFP-snail cells recruit a marginally greater percentage of neutrophils isolated from 4 out of 5 donors, compared to M2-GFP cell-derived CM. There was no difference in neutrophil migration induced by the CM harvested from M2-twist cells relative to CM from M2-GFP cells (Fig.2E). CM harvested from all three cells lines after TGF-β1/TNF-α treatment induced similar levels of neutrophil migration (Fig.2E), as did CM harvested from M2-twist KO cells treated with TGF-β1/TNF-α (Fig.2F). Next, we compared the neutrophil recruiting ability of CM derived from two aggressive breast cancer cell lines, M4 and BT549, and their twist KO counterparts. We found that CM from both twist KO cell lines retain robust neutrophil recruiting activity (Fig.2G-H). We did find that twist is required for the invasion ability of BT549 cells in a transwell invasion assay setup (Fig.S1G), suggesting that the neutrophil recruiting ability of these cells is not linked to their ability to invade. Similarly, there was no difference in the neutrophil recruiting ability of CM harvested from snail KD BT549 cells compared to CM derived from SCR control cells (Fig.2I). Together, these results provide evidence that snail or twist have no direct role of in regulating the ability of breast cancer cells to recruit neutrophils.

### TGF-β1/TNF-α treatment does not impact the neutrophil recruiting activity of TNBC cell lines

We previously reported that CM harvested from the TNBC cell lines M4, MDA-MB-231, and BT549 induces strong neutrophil migration (6). Here, we examined whether TGF-β1/TNF-α treatment further stimulates the neutrophil recruiting abilities of these cells. First, we characterized the effect of the dual treatment on EMT-associated markers. We measured an increased in Fn expression in both M4 and BT549 cells (Fig.S2Aiii,C). Dual treated M4 cells also exhibited greater expression of N-Cad (Fig.S2Aii), while the amount of E-Cad in M4 cells remained unchanged (Fig.S2Ai). In MDA-MB-231 cells, TGF-β1/TNF-α treatment marginally decreased expression of Fn (Fig.S2B). Finally, we found that the neutrophil recruiting activity of CM harvested from TNBC cells was unaffected by the TGF-β1/TNF-α dual treatment (Fig.S2D-F). Together, these results show that TGF-β1/TNF-α treatment elicits minor changes in the expression of EMT-associated markers in various TNBC cell lines and does not impact the neutrophil recruiting ability of TNBC-derived CM.

### TNF-α amplifies the neutrophil recruiting activity of M2 cells

TGF-β1 and TNF-α have regulatory roles during tumor progression (35, 52, 53). Both act synergistically in promoting EMT-associated changes in non-invasive breast cancer cells (36). To assess whether TGF-β1 and TNF-α synergize towards regulating neutrophil recruitment, we compared M2 cell responses to dual vs. single TGF-β1 or TNF-α treatments. We found that only dual-treated cells undergo dramatic morphological changes, losing epithelial sheet-like alignment and acquiring spindle-shaped mesenchymal features (Fig.3A). When analyzing EMT-associated markers, we found that while TNF-α failed to increase the expression of Fn and N-Cad, TGF-β1 upregulated the expression of both markers – a response that was further amplified with dual-treatment (Fig.3Bi,ii). Surprisingly, CM from TNF-α-treated M2 cells induced neutrophil migration as robustly as CM from dual-treated cells; besides, only a moderate increase in neutrophil migration occurred when using CM from TGF-β1-treated M2 cells (Fig.3C). These results indicate that TGF-β1 and TNF-α do not synergistically amplify the neutrophil recruiting activity of M2 cells. Next, we evaluated how the various treatments modulate the profiles of neutrophil chemokines secreted from M2 cells. We measured a 10-fold increase in the amount of CXCL1 in CM harvested from TNF-α-treated M2 cells compared to only a 4-fold increase when cells were dual-treated (Fig.3Di). We also noted a 2-fold increase in the amount of CXCL2 in the CM collected from TNF-α treated M2 cells (Fig.3Dii). In contrast, we detected a gradual increase in the amount of CXCL8 in the CM, with a 6-fold increase following TNF-α treatment and a 9-fold increase following dual-treatment (Fig.3Diii). While there was a 2-fold increase in the level of CXCL8 in response to TGF-β1, the later failed to induce any CXCL1 or CXCL2 from M2 cells (Fig.3Di,ii,iii). Together, these results suggest that the changes in neutrophil recruiting activity of M2 cells and alterations in EMT-associated markers are not interdependent. Furthermore, our findings reveal that treatment of M2 cells with TGF-β1 and/or TNF-α leads to changes in the CXCL1, CXCL2, and CXCL8 secretion profile and neutrophil recruitment ability of the CM.

**Figure 3.**
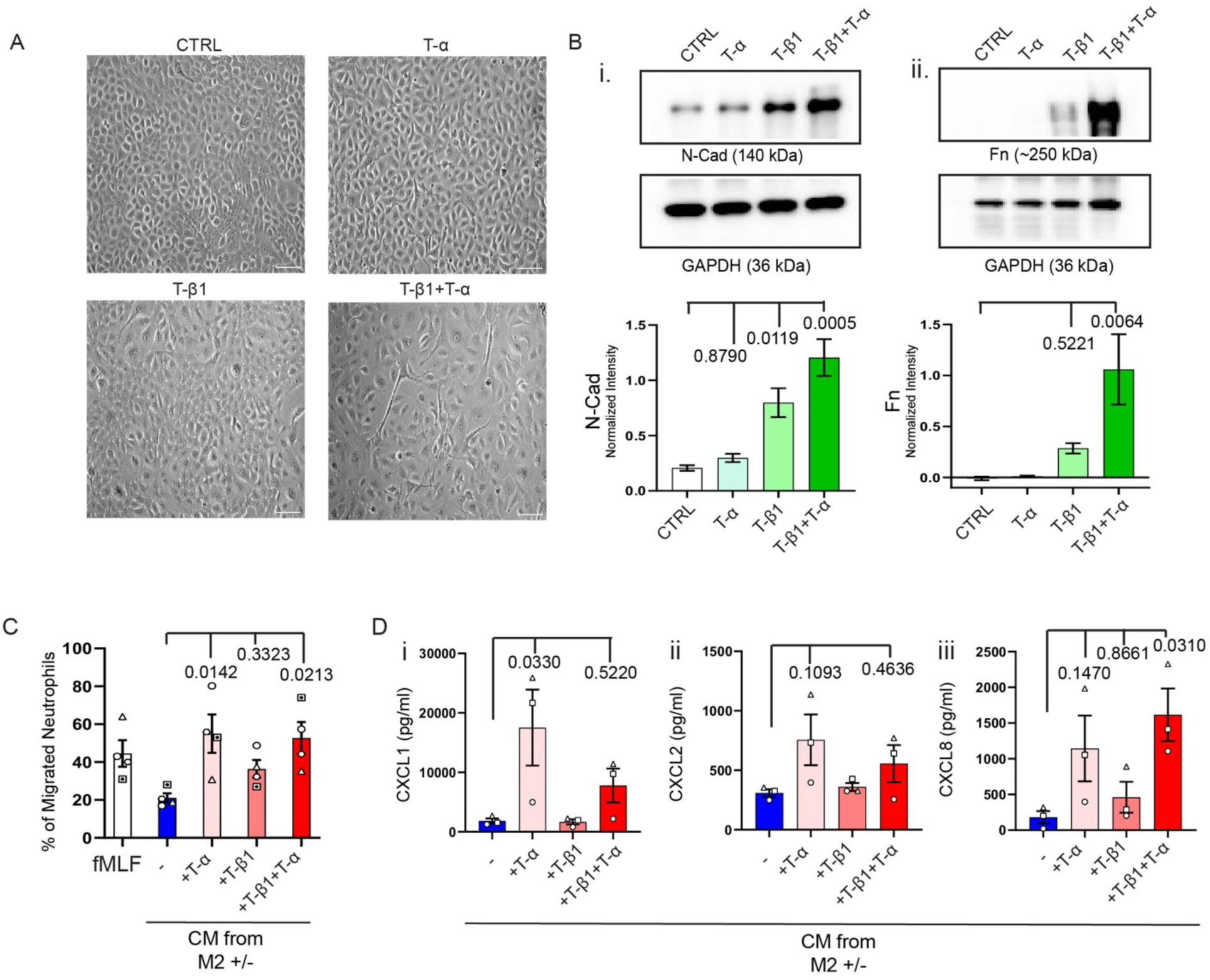
TNF-α treatment amplifies the neutrophil recruiting activity of M2 cells without inducing EMT associated changes. **A.** Representative (n=3) bright field images showing the morphology of CRTL, T-β1, T-α, or T-β1/T-α treated M2 cells. Scale bar=100 µm. **B.** Top: Representative western blots showing the expression of (i) N-Cad and (ii) Fn in CTRL and T-β1, T-α, or T-β1/T-α treated cells. Bottom: Graphs showing band intensities of the markers normalized to the respective loading controls (mean +/− SEM from n=3). **C.** Graph depicting the percentage of neutrophils that migrated into the bottom chamber of transwells containing equal volume of CM from CTRL, T-β1, T-α, or T-β1/T-α treated M2 cells (mean +/− SEM from n=4). Each dot represents response of neutrophils from independent donors. **D.** Graphs showing the amount of CXCL1 (i), CXCL2 (ii), and CXCL8 (iii) secreted from CTRL, T-β1, T-α, or T-β1/T-α treated M2 cells (mean +/− SEM from n=3). P values were determined using one-way ANOVA with Dunnett’s multiple comparisons test (B-D).

### p38MAPK regulates the secretion of neutrophil recruiting chemokines in TGF-β1/TNF-α treated M2 cells

To further understand the mechanisms underlying the amplified neutrophil recruiting activity of TGF-β1/TNF-α treated M2 cells, we examined key signaling pathways activated downstream of TGF-β1, TNF-α or TGF-β1/TNF-α dual-treatments. We first assessed the activation status of the TGF-β1-specific effector SMAD3 (54–56) and the TNF-α-specific effector NF-κβ (34, 57, 58) in M2 cells. As expected, we confirmed that TGF-β1 gives rise to SMAD3 phosphorylation (P-SMAD3) in M2 cells while TNF-α does not, and that the amount of P-SMAD3 remains similar in dual-treated vs. TGF-β1-treated M2 cells over periods of 1, 3, and 6 hrs after treatment (Fig.4A). We also confirmed that total SMAD3 levels are similar with both treatments (Fig. S3A). To assess the activity status of NF-κβ, we monitored the nuclear translocation of p65, a component of the p50/p65 heterodimer (59). We found that TNF-α treatment induces a strong nuclear translocation of p65 in M2 cells – seen and quantified as a high nuclear/cytosolic p65 ratio (Fig.4Bii,v). In contrast, p65 remained cytosolic in TGF-β1-treated M2 cells and, as expected, in control cells (Fig.4Bi,iii,v). Interestingly, we detected a significant increase in the nuclear/cytosolic p65 ratio when M2 cells are dual-treated with TGF-β1/TNF-α compared to a single TNF-α treatment, suggesting cross-talk between the two pathways (Fig.4Biv,v). Next, we evaluated the activation of MAPK and AKT signaling pathways in dual-treated M2 cells, as these downstream signaling events are known to promote EMT-associated changes in cancer cell lines (20, 36, 60). Over periods of 1, 3, and 6 hrs after treatment, we measured a strong and sustained p38MAPK phosphorylation (P-p38MAPK) signal in response to TGF-β1/TNF-α dual-treatment in M2 cells (Fig.4Ci), while total p38MAPK levels were similar irrespective of treatment (Fig.S3B). In contrast, no change in the phosphorylation amount or total amount of either ERK, JNK or AKT with TGF-β1/TNF-α treatment was measured (Fig.4C ii-iv, Fig.S3C-E).

**Figure 4.**
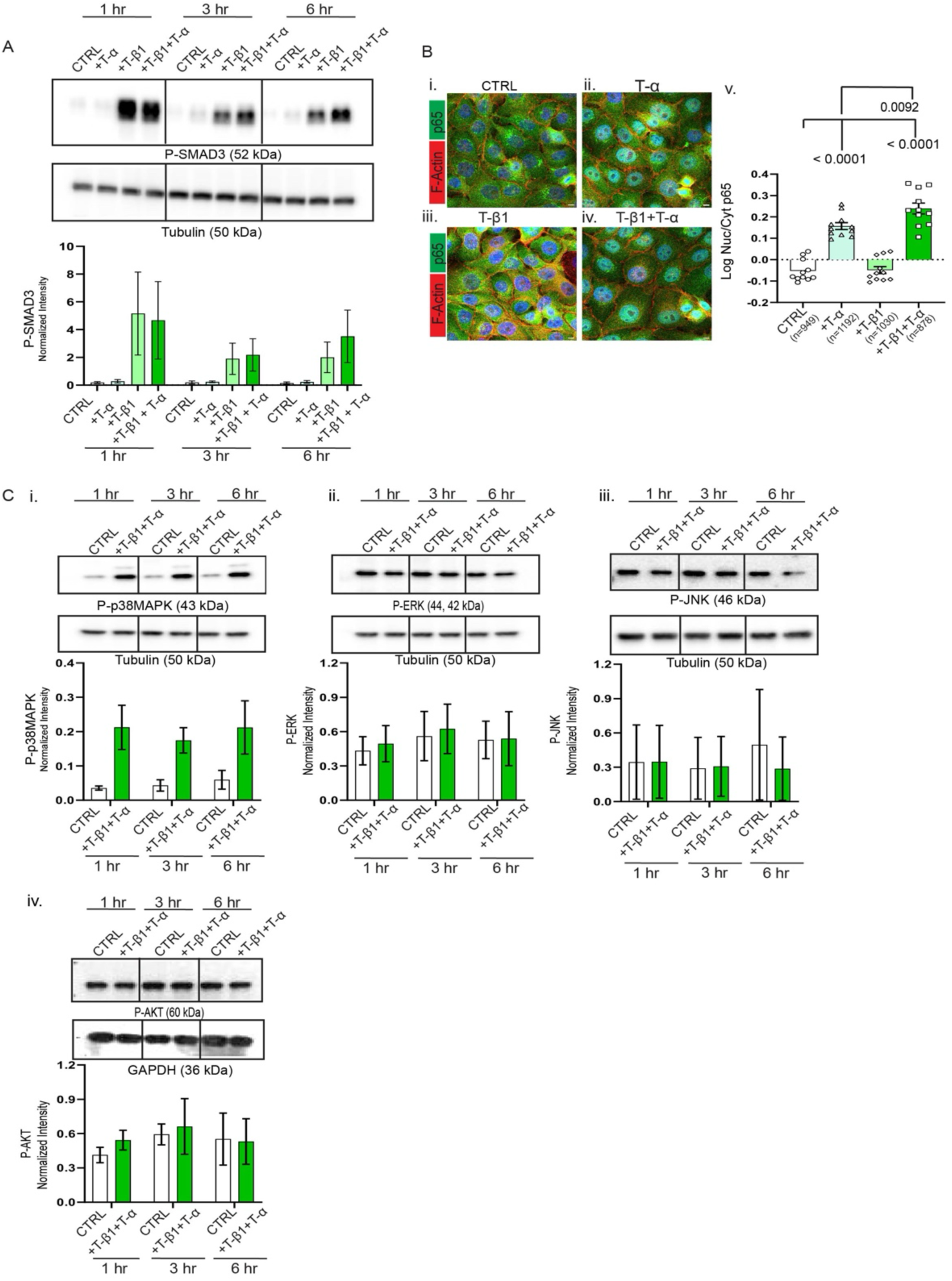
Signaling pathways activated in M2 cells treated with TGF-β1/TNF-α. **A.** Top: Representative western blots showing P-Smad3-specific bands over the course of 1, 3, and 6 hrs of treatments. Bottom: Graphs depicting band intensities of P-Smad3 normalized to the loading control (mean +/− SEM from n=3). Western blot of total Smad3 is provided in Fig. S3. **B.** Representative IF images showing MIPs of fixed CRTL M2 cells or M2 cells treated for 30 min with T-β1, T-α, or T-β1/T-α and stained for p65 (green), F-actin with phalloidin-TRITC (red), and nucleus (blue) (i-iv). Scale bar =5 µm. Graph (v) depicting the nuclear/cytosolic intensity ration for p65. Each dot represents average of all cells in each image (3 images/condition/experiment). Total no. of cells (n) analyzed is depicted under each condition. **C.** Representative western blots showing P-p38MAPK (i), P-ERK (ii), P-JNK (iii) or P-AKT (iv) specific bands over the course of 1, 3, and 6 hrs of treatments. Graphs show band intensities of the respective phospho-proteins normalized to the loading control (mean values +/− SEM from n=3). Western blots of total p38MAKP, ERK, JNK, and AKT are provided in Fig. S3. P values were obtained using one-way ANOVA with Dunnett’s multiple comparisons test (Bv).

Next, we dissected the effect of TGF-β1 and TNF-α on p38MAPK activation. While individual TGF-β1 or TNF-α treatments induced moderate P-p38MAPK without affecting total protein levels, dual-treated M2 cells exhibited a significant increase in P-p38MAPK relative to control cells at 1 and 6 hrs post stimulation (Fig.5A, Fig.S3F). As p38MAPK activity is known to regulate the expression of EMT-associated markers in breast epithelial and cancer cell lines (61) and also controls the expression of several cytokines and chemokines (62–65), we next assessed the effect of p38MAPK inhibition on TGF-β1/TNF-α-mediated EMT markers and chemokine production in M2 cells. We analyzed the morphology of co-treated M2 cells in the presence of vehicle control or doramapimod (DMPM), a potent p38MAPK inhibitor (66, 67), at a dose that effectively inhibits P-p38MAPK (Fig.5B) but maintains total p38MAPK levels (Fig.S3G). Interestingly, we found that the mesenchymal phenotypes induced by dual-treatment is retained in the presence of the inhibitor (Fig.5C). While we measured a small decrease in the expression of Fn with the inhibitor treatment (Fig.5C;Diii), we observed no changes in the upregulation of N-Cad and Vim expression in co-treated M2 cells in the presence of the inhibitor (Fig.5C;Di-ii). In contrast, we measured a substantial decrease in the amount of both CXCL1 and CXCL8 in the CM harvested from dual-treated M2 cells in the presence of the inhibitor compared to the vehicle control (Fig.5Ei,ii). Together, these findings indicate that p38MAPK activity occurring downstream of TGF-β1/TNF-α dual-treatment regulates the expression of neutrophil recruiting chemokines independently of EMT-associated morphological and markers changes. Further, these findings add to the observation that EMT can be uncoupled from the secretion of neutrophil chemokines in M2 cells.

**Figure 5.**
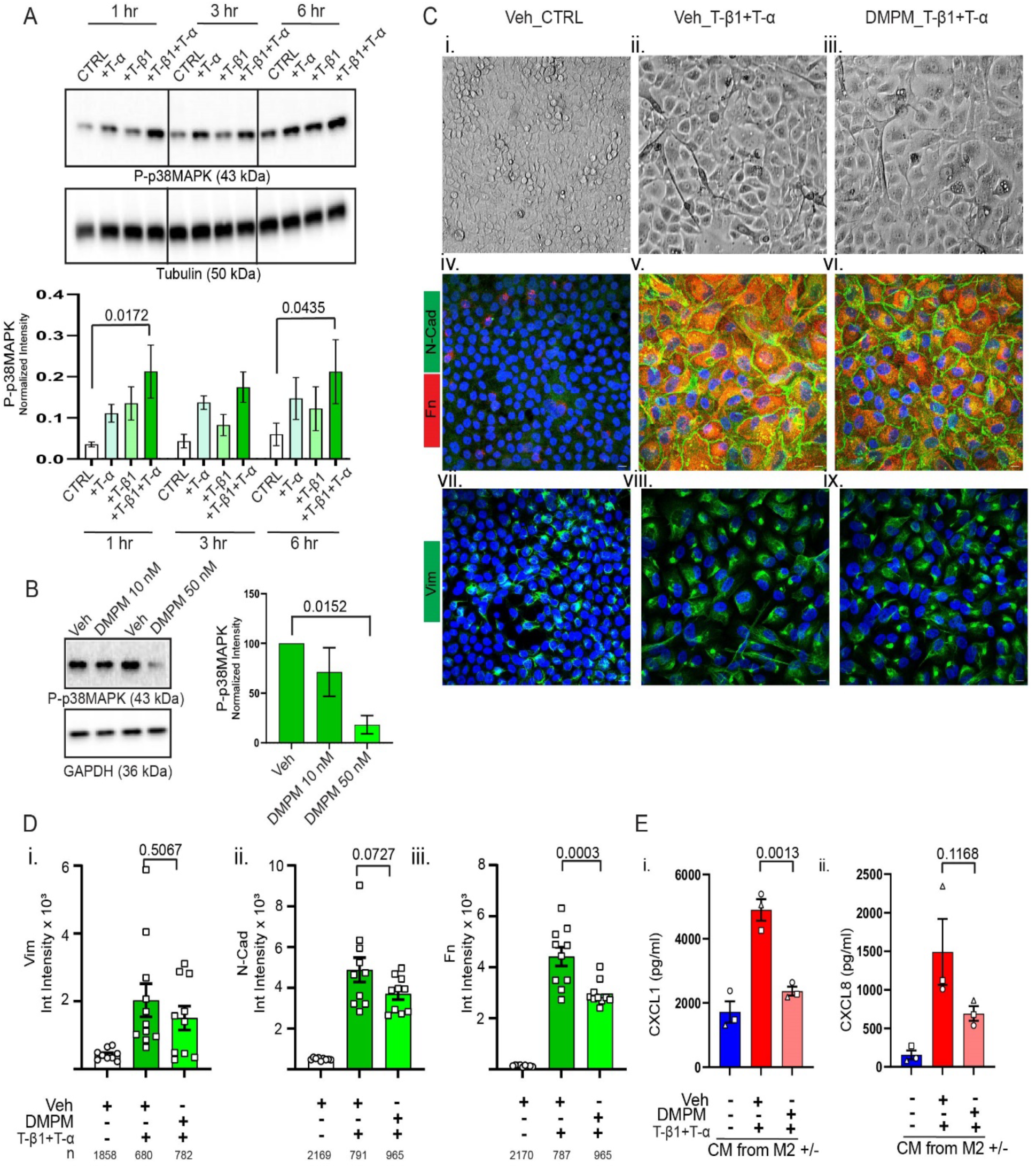
P-p38MAPK regulates the secretion of neutrophil recruiting chemokines in TGF-β1/TNF-α treated M2 cells. **A.** Top: Representative western blot showing P-p38MAPK-specific bands in CTRL M2 cells or M2 cells treated with T-β1, T-α or T-β1+T-α over the course of 1, 3, and 6 hrs. Bottom: Graph depicting band intensities of P-p38MAPK normalized to the loading control (mean +/− SEM from n=3). Representative western blot of total p38MAKP is provided in Fig. S3. **B.** Left: Representative western blot showing P-p38MAPK-specific bands in M2 cells pre-treated with 10nM or 50nM of DMPM or vehicle control and stimulated with T-β1+T-α for 72 hrs. Right: Graph depicting band intensities of P-p38MAPK normalized to loading control and represented as percentage of vehicle control (mean +/− SEM from n=3). Representative western blot of total p38MAKP is provided in Fig. S3. **C.** Representative bright filed (i-iii) or IF (iv-ix) images (n=3) showing MIPs of M2 cells pre-treated with DMPM or vehicle control and stimulated with T-β1+T-α for 72 hrs. Cells were stained for N-Cad (green)/Fn (red) (iv-vi) or Vim (green) (vii-ix) and nucleus (blue). Scale bar 20 µm (bright field) or 10 µm (IF). **D.** Graphs depicting integrated intensity measures of Vim (i), N-Cad (ii), and Fn (iii). Each dot represents average of all cells in each image (≥3 images/condition/experiment). Total no. of cells (n) analyzed is depicted under each condition. **E.** Graphs showing amount of CXCL1 (i) and CXCL8 (ii) secreted by T-β1+T-α treated M2 cells in the presence of DMPM or vehicle control (mean values +/− SEM from n=3). P values were determined using two-way (A) and one-way (B, D, E) ANOVA with Dunnett’s multiple comparisons test.

### Analysis of RNA sequencing datasets from breast cancer cell lines identifies correlations between *CXCL1, CXCL8,* TGF-β and TNF-α pathways

To expand our findings to additional cancer cell lines *ex vivo* we analyzed two complementary datasets: (i) the Harvard Medical School LINCS Breast Cancer Profiling Project (HMS LINCs dataset), which includes 35 breast cancer cell lines of known tumor subtype, and (ii) the Broad Institute Cancer Dependency Map Project (DepMap), containing a dataset of gene expression profiles for 1450 cancer cell lines of varying lineages including 68 breast cancer cell lines.

First, we tracked *CXCL8* and *CXCL1* expression across different breast cancer molecular subtypes using the HMS LINCS dataset (Fig.6A). We measured a significant difference in *CXCL8* and *CXCL1* mRNA transcripts in TNBC cell lines, compared to HR^+^ cell lines (Fig.6B,C). We then tested whether this increase is due to a correlative relationship between *CXCL1* and *CXCL8* by calculating the correlation coefficient (reported as Pearson r) of all 29,187 surveyed genes against *CXCL8* across the 35 available cell lines. Supporting our findings in M2 cells, we found that the expression of *CXCL8* correlates with *CXCL1* (R^2^=0.33’), *CXCL2* (R^2^=0.65) and *CXCL3* (R^2^= 0.53) (Fig.6D,E). We also found that a large set of genes belonging to the TGF-β signaling pathway is positively correlated with *CXCL8* (Fig.6F), including transforming growth factor beta induced (*TGFΒI*, R^2^=0.55) (Fig.6G), *TGFΒR2* (R^2^=0.54) and, to a lesser extent, *SMAD3* (R^2^=0.31). Similarly, we identified members of the TNF-α signaling pathway to significantly correlate with *CXCL8* (Fig.6F), including *BIRC3* (R^2^=0.63) (Fig.6H), a TRAF1/2 binding, anti-apoptotic protein that is upregulated with breast cancer metastasis (68–70), TNF-α induced protein 3 (*TNFAIP3*, R^2^=0.55), known to regulate NFκB and protect breast cancer cells from TNF-α-induced cell death (71), and *CD137* (*TNFRSF9*, R^2^=0.46), which is associated with breast cancer metastasis (72) (the full list of *CXCL8*-correlating genes is given in Table S2). Together, the HMS LINCS dataset analysis validates our experimental findings by identifying an association between *CXCL8* and *CXCL1* and key effectors of TGF-β and TNF-α signaling.

**Figure 6.**
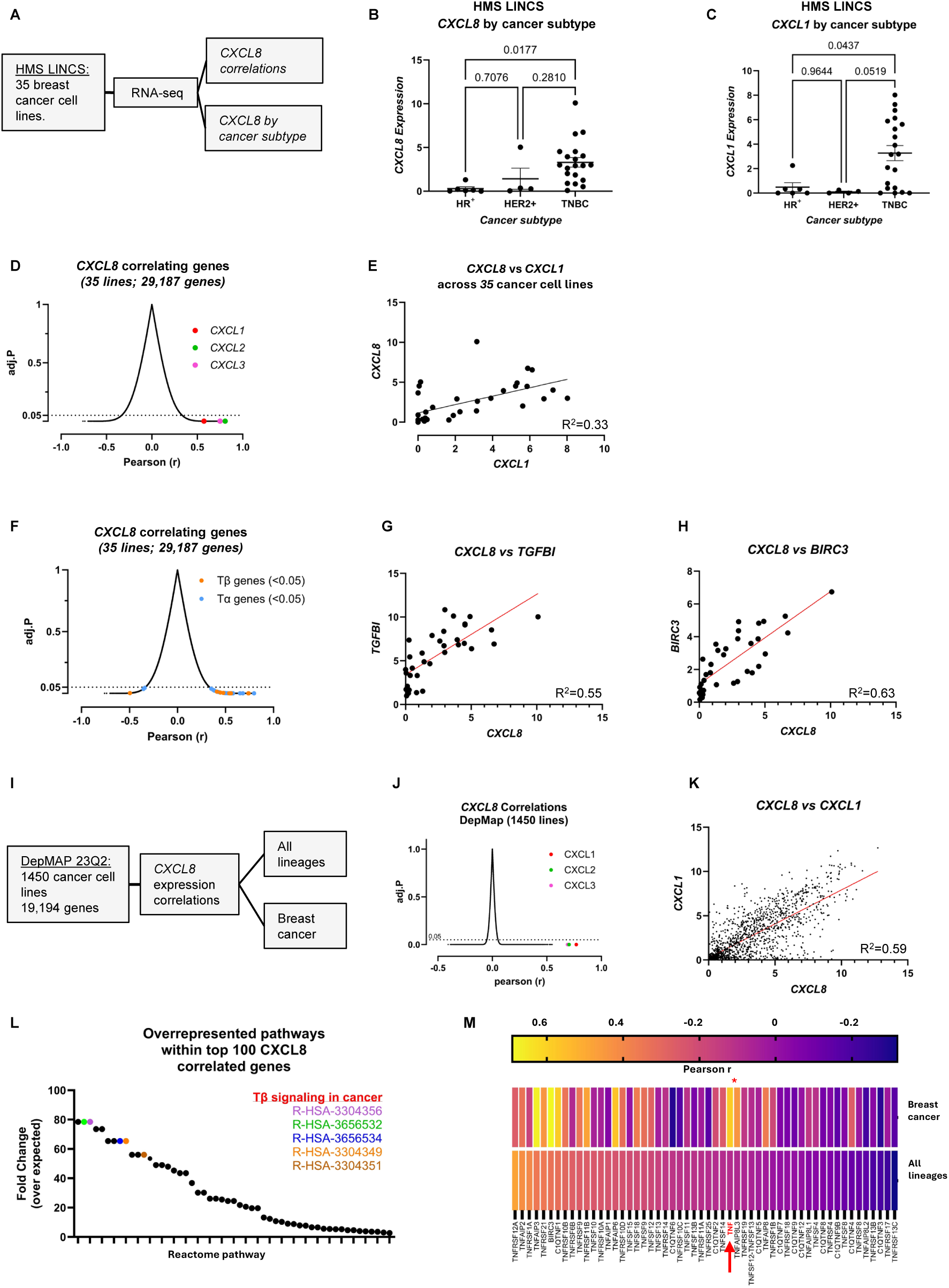
Computational analysis of gene expression datasets identifies the association between CXCL8 and members of TGF-β1/TNF-α pathways. **A.** Workflow for the analysis of RNAseq dataset from the Harvard Medical School Breast Cancer Profiling Project (ID: 20352). **B-C.** Expression level of CXCL8 (B) and CXCL1 (C) in HR+, HER2+, or TNBC cell lines available in the LINCs dataset **D-H.** Distribution of linear correlations (Pearson r) of all genes surveyed in the LINCs dataset against CXCL8 (D,F). Individual gene correlation between CXCL8 and CXCL1 (E), CXCL8 and TGFBI (TGFBI transforming growth factor beta induced) (G) and CXCL8 and BIRC3 (H) across the 35 cell lines. **I.** Workflow for the analysis of Broad institute DepMap gene expression profiles. **J-K.** Distribution of linear correlations (Person r) between CXCL8 and all genes surveyed in the DepMap dataset, across 1450 available cell lines (J). Correlation between CXCL1 and CXCL8 across available cell lines (K). **L.** Analysis of the top 100 CXCL8 correlation genes using panther overrepresentation assay. T-β related pathways are marked in color and their Reactome pathway ID provided. **M.** DepMap Analysis of CXCL8 correlations against T-α genes in breast cancer lines vs all lineages. The heatmap presents correlation coefficients of (Pearson r). TNF gene is highlighted in red.

We next analyzed gene expression data from DepMap (Fig.6I). In agreement with our experimental data and the HMS LINCS analysis, we found that *CXCL8* expression is strongly correlated with *CXCL1* (R^2^=0.59), followed by *CXCL2* (R^2^=0.49) and *CXCL3* (R^2^=0.48), across all DepMap cancer cell lines (Fig.6J,K). We then analyzed the top *CXCL8* correlating genes for statistical over-representation of signaling pathways using the Panther and Reactome datasets (73, 74). We found that TGF-β related genes show high fold-enrichment within the top 100 *CXCL8* correlated genes across all cancer cell lines. Specifically, we noted a 78-fold increase above expected in “*Reactome pathway SMAD2/3 Phosphorylation Motif Mutants in Cancer*” (R-HSA-3304356), and the related “*TGFΒR1 KD Mutants in Cancer*” (R-HSA-3656532), “*Loss of Function of TGFΒR1 in Cancer*” (65-fold, R-HSA-3656534), “*Loss of Function of SMAD2/3 in Cancer*” (65-fold, R-HSA-3304349), and “*TGF-β Receptor Complex in Cancer*” (56-fold, R-HSA-3304351) (respectively highlighted in magenta, green, blue, orange and brown in Fig.6L). Interestingly, no TNF-α related genes were overrepresented in the top *CXCL8-*correlated gene from DepMap. This contrasted with the HMS LINCS dataset, where only breast cancer cell lines are included. Next, we tested whether correlations with TNF-α pathway genes were stronger in breast cancer cell lines relative to all cell lines tested in DepMap. We observed that most genes correlated similarly in all cell lines vs. breast cancer cell lines (R^2^=0.86), including TGF-β-related genes. Interestingly, however, we identified many TNF-α pathway genes to associate more strongly with *CXCL8* when comparing breast cancer cell lines to the entire dataset. Specifically, we observed significant associations between *CXCL8* and TNF-α genes (Fig.6M) in breast cancer cell lines. Besides, we measured an increased association between *CXCL8* and expression of *TNF* itself in breast cancer cell lines (R^2^=0.34) vs. all cell lines (R^2^=0.01). The increased correlation with *CXCL8* expression was also observed for genes found in the HMS LINCS dataset, including *TNFAIP3* (R^2^=0.45 vs. R^2^=0.12 in all cell lines), *BIRC3* (R^2^=0.47 vs. R^2^=0.11), and to a much lesser extent, *TNFRSF9* (R^2^=0.11 *vs.* R^2^=0.07). These findings complement and provide a broader context for the interpretation of our findings when analyzing the HMS LINCS dataset, and further suggests a regulatory interaction between TGF-β, TNF-α and *CXCL8* in breast cancer.

Finally, we tested whether *CXCL8* and *CXCL1* expressions correlate with EMT associated genes including *VIM* (gene for Vim), *SNAI1* (gene for Snail1), and *TWIST1* (gene for Twist1) in breast cancer cell lines using the HMS LINCS dataset. As expected, we measured a significantly higher *VIM* expression in the TNBC subtypes compared to others (Fig.S4A) (12). However, we observed low correlations for both *CXCL8* (R^2^=0.11) and *CXCL1* (R^2^=0.01) expressions with *VIM* in the TNBCs (Fig.S4B). Similarly, we measured weak correlations for both *CXCL8* and *CXCL1* expressions with *SNAI1* (R^2^=0.01 *CXCL8*, R^2^=0.01 *CXCL1*) and *TWIST1* (R^2^=0.03 *CXCL8*, R^2^=0.10 *CXCL1*) (Fig.S4C,D) in the TNBCs. When we expanded our analysis to the DepMap dataset, we again observed weak to no correlations between *VIM* and *CXCL8* (R^2^=0.01) or *CXCL1* (R^2^=0.00) (Fig.S4E,F). In contrast to the chemokines, VIM expression showed higher correlation with *CDH2* (gene for N-Cad) (R^2^=0.22) (Fig.S4G). When we focused on the 68 breast cancer cell lines from the DepMap dataset we measured increased correlation of *VIM* expression with *CDH2* (R^2^=0.5) (Fig.S4J). However, as seen with the HMS dataset, we observed low correlations between VIM and either *CXCL8* (R^2^=0.11) or *CXCL1* (R^2^=0.13) (Fig.S4H,I). These computational analyses further substantiate the lack of an association between the regulation of *CXCL8* and *CXCL1* expression and EMT-associated changes in breast cancer cells, and confirm the existence of a TGF-β/TNF-α regulatory axis for *CXCL1* and *CXCL8* that is, again, independent of EMT induction in breast cancer cells.

## DISCUSSION

In this study we aimed to address whether TGF-β and TNF-α, two pro-inflammatory factors frequently detected in tumor niches, impact neutrophil recruitment to breast tumors. We found that dual treatment with TGF-β1 and TNF-α enhances the secretion profile of neutrophil chemokines from human breast epithelial cell lines in a manner that is independent of the parallel induction in EMT markers. We also discovered that, unlike the expression of EMT markers, the TGF-β1/TNF-α-dependent upregulation of the neutrophil chemokines CXCL1 and CXCL8 relies on activation of the p38MAPK pathway. Finally, by analyzing transcriptomic databases for a wide array of human cancer cell lines representing different types of solid tumors including breast, we uncovered significant correlations between *CXCL8* and a number of TGF-β1/TNF-α associated genes. Further, these computational analyses did not support a link between *CXCL8* or *CXCL1* and EMT-associated genes, thus broadening the significance of our studies involving the M2 breast epithelial cell line.

EMT-associated changes transform cancer cells into more aggressive and invasive types. The presence of immune cells such as tumor-associated macrophages and other myeloid cells has been linked to expression of EMT markers in various cancer types (7). Here, we confirmed the occurrence of dramatic changes in EMT marker expression in M2 cells treated with TGF-β1/TNF-α. Treated M2 cells also acquired robust neutrophil recruitment activity. However, our study provides five lines of evidence showing that such EMT-associated changes are not essential for neutrophil recruitment. First, the ability of M2 cells to recruit neutrophils is greatly amplified with TNF-α treatment even though the latter fails to bring EMT-associated changes. Second, ectopic expression or depletion of snail and twist in M2 cells does not perturb the neutrophil recruitment ability of the CM obtained from these cells. Furthermore, depleting these TFs from aggressive breast cancer cell lines, e.g. M4 and BT549, has no effect on their robust neutrophil recruitment activity. Third, the neutrophil recruitment activity of aggressive breast cancer cell lines, e.g. M4, MDA-MB-231, and BT549, is not further amplified by TGF-β1/TNF-α treatment, even though the latter enhances expression of key EMT markers. Fourth, blocking p38MAPK activation drastically reduces the amount of neutrophil guidance cues in the CM obtained from TGF-β1/TNF-α dual-treated M2 cells and yet, mesenchymal morphology and levels of EMT markers are maintained under these conditions (except for Fn, which showed a partial decrease). Fifth, transcriptomic analysis of large datasets of cancer cell lines of different lineages including breast cancer provided additional evidence of a lack of correlation between neutrophil guidance cues (*CXCL8, CXCL1*) and EMT-associated genes. Together, these findings indicate that EMT-associated changes and amplified neutrophil recruitment are two distinct outcomes of TGF-β1/TNF-α stimulation.

The amplified neutrophil recruitment ability of treated M2 cells was accompanied by increases in secreted CXCL1, CXCL2, and CXCL8. CXCL8 is a potent ligand for CXCR1 and CXCR2, both of which are expressed in human neutrophils (75, 76), whereas CXCL1 and CXCL2, which are members of the GRO family of chemokines, are CXCR2-specific ligands. All three chemokines have well-established role in mediating neutrophil recruitment to tumors (77–79), and the increased expression of all three chemokines is associated with more aggressive tumors, including glioblastoma, breast and colorectal cancer, and malignant melanoma (75, 80–82). We also found that expression of *CXCL8* mRNA is significantly higher in aggressive TNBC cell lines compared to HR+ ones. Furthermore, we discovered that the expression of *CXCL8* is highly correlated with that of *CXCL1/2* transcripts in breast cancer cell lines. We previously described a key role for CXCL1/2 chemokines secreted by aggressive breast cancer cell lines to induce robust neutrophil recruitment (6). In the current study, we found that neutralizing CXCL8 action did not completely abrogate the ability of CM from TGF-β1/TNF-α treated M2 cells to induce a neutrophil chemotactic response. Therefore, CXCL1/2 and CXCL8 may act in a synergistic manner to control neutrophil trafficking to breast tumors.

While the role of TGF-β1/TNF-α in regulating the expression of EMT markers is well-established, how the combined action of the two factors impacts immune cell recruitment to tumors has been largely unexplored. TNF-α induces the expression and secretion of CXCL1, CXCL10 and several CCL chemokines in immune cells, endothelial cells and cancer cells (83–85). TGF-β controls chemokine availability either by inducing or suppressing the expression and secretion of chemokines (86–88). For instance, TGF-β promotes neutrophil recruitment by upregulating the expression and secretion of CXCL5 in hepatocellular carcinoma (86) while it suppresses the secretion of CXCL1 by mesenchymal stromal cells (87). Because we previously reported that CXCL5, CCL2, CCL3, and CCL5 occur at low levels in CM collected from aggressive breast cancer cell lines, here we focused on other abundantly secreted chemokines (6). When we dissected the effect of TGF-β1 and TNF-α through individual vs. dual treatment, we found interesting changes in the chemokine secretion profiles of M2 cells. Treatment with TNF-α was effective at inducing CXCL1, CXCL2, and CXCL8, while TGF-β1 was not. Treatment with TGF-β1 suppressed CXCL1 secretion from the co-treated M2 cells, supporting previous findings (87). On the other hand, the amount of secreted CXCL8 was higher in cells treated with both TNF-α and TGF-β1 compared to cells stimulated with TNF-α alone, suggesting crosstalk between the two signaling pathways. Additionally, the positive correlation we measured between CXCL8 and a number of TGF-β1 and TNF-α-associated genes in a large dataset of breast cancer cell lines validates our *in vitro* findings on a positive regulation of CXCL8 by TGF-β1 and TNF-α. Our findings also suggest that the relative availability of TGF-β1 and TNF-α in the tumor niche controls the abundance of key neutrophil recruiting chemokines and thus regulates the degree of neutrophil recruitment to breast tumors.

To seek mechanistic insight, we evaluated the status of signaling pathways known to impact EMT programs, including Smad3, NF-κβ, p38MAPK, ERK, JNK, and AKT (20). Of all the pathways tested, we found more robust activation of NF-κβ and p38MAPK pathways with dual TGF-β1/TNF-α treatment compared to single treatments. Yet, we found a key and specific role for p38MAPK-mediated signaling in neutrophil recruitment towards CM isolated from TGF-β1/TNF-α treated M2 cells. Active p38MAPK is already known to regulate the production of a number of pro-inflammatory cytokines and chemokines from both immune cells and non-immune cells, such as epithelial and endothelial cells (62–65). Such regulation can occur through the modulation of NF-κB activation or by promoting mRNA stability (89, 90). Furthermore, a regulatory effect of p38MAPK on chemokines and Fn expression has been reported in breast cancer (91).

Our studies shed additional light into this. Indeed, we found that, while Vim and N-Cad expression levels remained unchanged, the presence of a pan-P38MAPK inhibitor (67) resulted in reduced secretion of CXCL1 and CXCL8 as well as a decrease in Fn expression. Other pro-inflammatory factors, e.g., IL-1β, have been shown to regulate Fn expression (92). Our observation of a partial reduction in Fn following inhibitor treatment suggests that Fn can be regulated by EMT-independent mechanisms. Future studies should determine which isoform/s of p38MAPK are specifically involved in regulating the secretion of neutrophil guidance cues with the TGF-β1/TNF-α treatment and elucidate the mechanisms underlying the regulation.

Neutrophil recruitment to tumors is associated with breast cancer progression. The findings we report here establish that neutrophil recruitment can occur independently of EMT in breast tumors, and that the TGF-β1/TNF-α /p38MAPK signaling axis is a potential predictor of neutrophil recruitment in the breast tumor niche. Future studies will aim at understanding how TGF-β1/TNF-α exposure impacts neutrophil function and influences breast cancer progression.

## MATERIALS AND METHODS

### Materials

TGF-β1 from R&D Systems (7754-BH-005) or Peprotech (100-21); TNF-α from Peprotech (300-01A) or Prospec (cyt-114); human recombinant CXCL8 from ProSpec (CHM349); Formyl-methionyl-leucyl-phenylalanine (fMLF) from Sigma-Aldrich (F3506); DMSO from Sigma-Aldrich; Doramapimod from Cayman Chemicals (10460), human CXCL-8 antibody from R&D systems (MAB208-100), and isotype antibody from Invitrogen (02-6100 or 31903) were used in the study.

### Isolation of Human Neutrophils

Blood was obtained from healthy human male and female subjects aged 19–65 years, through the Platelet Pharmacology and Physiology Core at the University of Michigan. The Core maintains a blanket IRB for basic science studies, where HIPAA information is not required. Therefore, while the IRB-approved Core enrolls healthy subjects that conform to the protection of human subject standards, we did not have access to this information. The samples that we received were fully de-identified. Neutrophils were purified using dextran-based sedimentation followed by histopaque-based density gradient centrifugation as described previously (6). Cells were more than 99% viable immediately following isolation. To address donor-to-donor variability of neutrophil response, cells were routinely tested for minimum basal activity and a robust response to fMLF stimulation as described before (6).

### Cell lines

We used a panel of 10 cell lines of human breast epithelial origin described in Table 1. MCF10A (M1), MCF10AT (M2), and MCF10CA1a (M4) were obtained from the Karmanos Research Institute. The human triple-negative breast cancer (TNBC) cell line BT549 and MDA-MB-231 were purchased from ATCC. M2-GFP, M2-snail, M2-twist, M2 twist KO, M4 twist KO, and BT549 twist KO cell lines were generated and validated in the lab for the expression of snail, twist, and GFP (see Fig.S1). All cell lines were maintained using representative culture media (Table 1) and incubated at 37°C in 5% CO_2_ humidified tissue culture incubator. All cell lines tested negative for mycoplasma contamination using the Mycoalert detection kit (Lonza). All cell lines were verified and authenticated based on short tandem repeat (STR) markers (Biomedical Research Core Facilities).

**Table 1.**
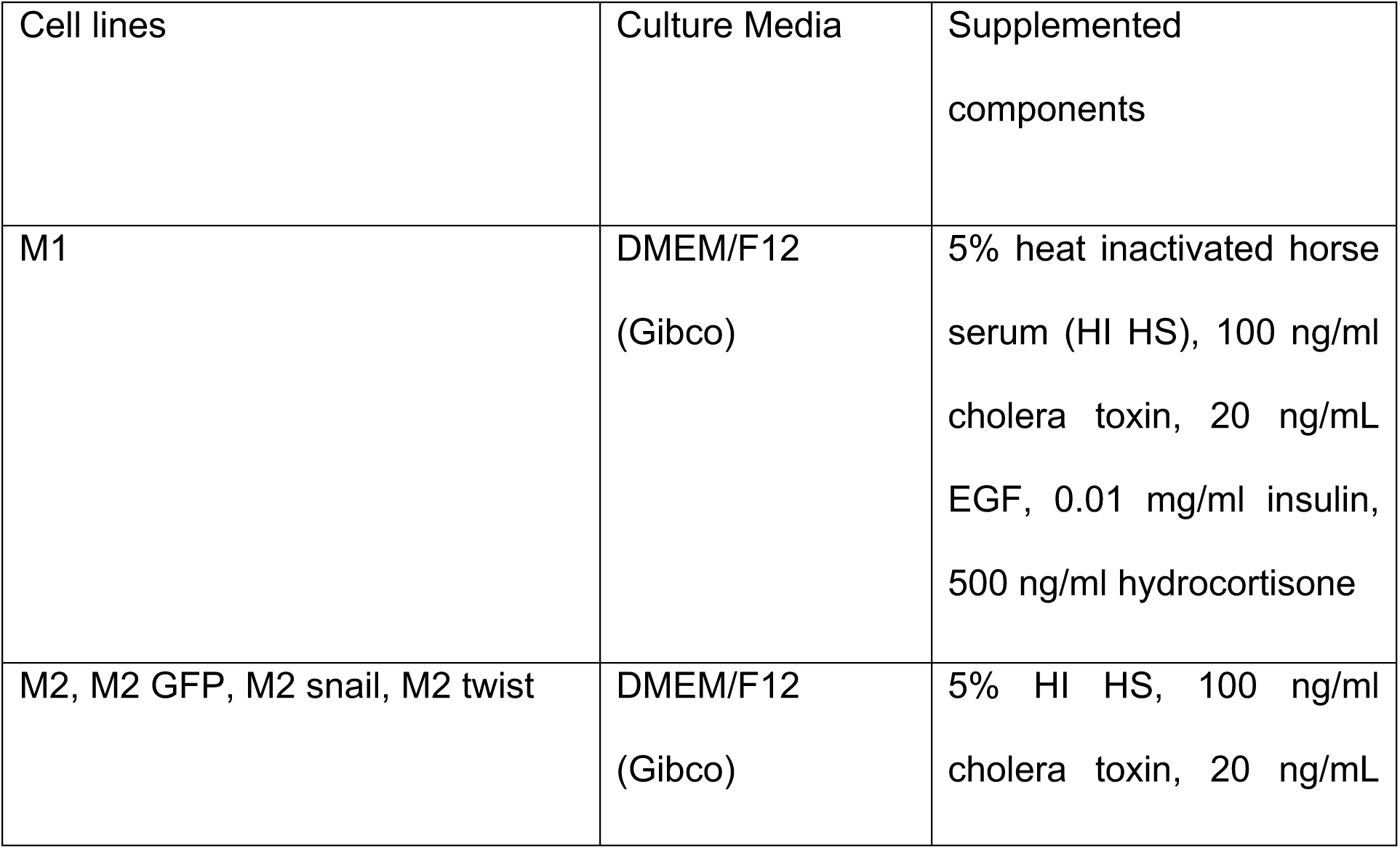
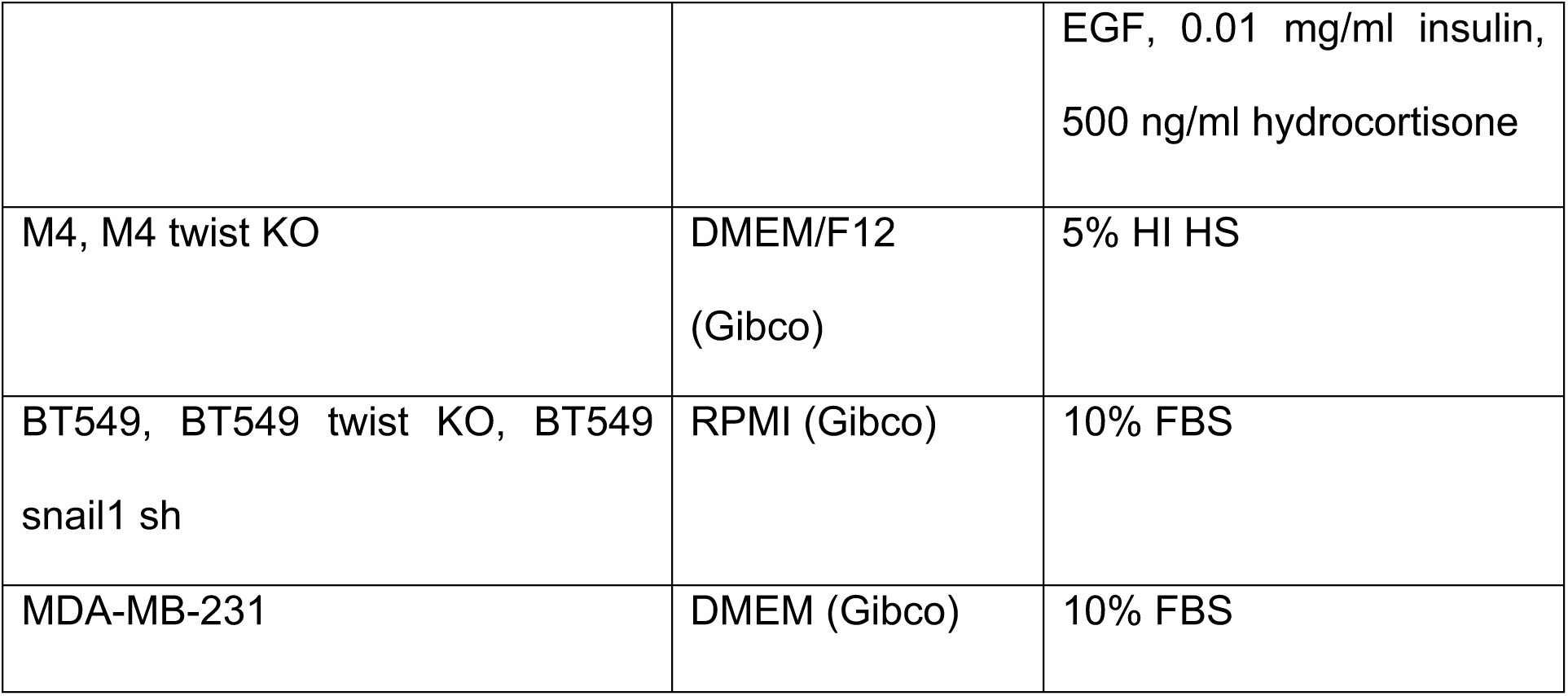
Cell lines and culture media used on the study.

### Plasmid constructs

The pBABE-Puro-twist, -eGFP-snail6SA, or -GFP constructs were used to generate stable M2 cell lines expressing twist, eGFP-snail6SA or GFP control. Twist from pWZL-Blast-twist construct (a kind gift from Jing Yang, UCSD) or eGFP-snail6SA from the eGFP-Snail6SA construct [Addgene plasmid #16228, deposited by Mien-Chie Hung; (48)] was subcloned into pBABE-Puro backbone from the pTK92 [Addgene plasmid #46356, deposited by Iain Cheeseman; (93)]. pBABE-GFP was generated from pTK92. For CRISPR-based knockouts, twist1 sgRNA CGGGAGTCCGCAGTCTTACGAGG was cloned into pLentiCRISPRv2 construct as described [Addgene plasmid #52961, deposited by Feng Zhang, (94)]. The pGipZ V3LHS_328731 and pGipZ scramble (SCR) constructs, obtained from the Vector core at the University of Michigan, were used to generate stable BT549 cell lines expressing snail1 and SCR shRNA, respectively.

### Stable cell line generation

For generating M2 cell lines stably expressing exogenous proteins, pBABE-based constructs were transfected using Lipofectamine^TM^ 3000 (Invitrogen) into Phoenix 293T packaging cells. Retroviral particles released from the packaging cells were collected 48 hrs post-transfection and concentrated using PEG-8000 (V3011, Promega), and then used to infect M2 cells in the presence of 10 µg/ml polybrene. The clones expressing the constructs were then selected in 2 µg/ml puromycin and verified using western blotting and fluorescence microcopy. For generating CRISPR-based knockout (KO) or shRNA-based knockdown (KD) cell lines, pLentiCRISPRv2 construct carrying twist1 sgRNA or pGipZ construct carrying snail1/SCR shRNA was produced by the Vector Core at the University of Michigan and were infected as described above. Clones expressing the construct were selected with puromycin and verified by western blotting and sequencing.

### Treatment for Harvesting Conditioned Media

Cell lines were seeded at a density of 0. 03×10^6^/ml in 2 ml volume of complete medium in each well of a six well tissue culture plate and incubated overnight to allow cell adhesion. The following day, the cells were either left untreated, or treated with 20 ng/ml TGF-β1 and 100 ng/ml TNF-α for 72 hrs at 37°C. For inhibiting p38MAPK phosphorylation, cells were pre-treated with doramapimod (DMPM) or vehicle control for 2 hrs followed by TGF-β1/TNF-α treatment. The culture medium was then removed, and cells were gently washed twice with calcium and magnesium free sterile DPBS (Gibco) to remove left over serum containing media. Cells were then incubated with fresh medium without serum and incubated for an additional 48 hrs. The media were harvested and filtered through 0.22 μm membrane filters to remove dead cell debris. Aliquots were frozen at −30°C until analyzed.

### Harvesting Conditioned Media from WT vs. KO or KD Cancer Cell Lines

To generate conditioned media (CM), breast cancer cell lines were seeded at a density of 0.15×10^6^/ml in 2 ml (6-well tissue culture plate) of complete medium for 24 hrs, at which point they reached ~ 70% confluence. The culture medium was then removed, and cells were gently washed as mentioned above and incubated with serum free fresh medium for an additional 48 hrs. The CM was collected and stored as described above.

### Bright-Field Microscopy

To assess morphological changes with treatment, cell lines were treated as before for 72 hrs. Bright-field images in three randomly selected fields per condition were captured using a 10x objective lens on a Zeiss Axiovert microscope.

### Immunofluorescence Microscopy

For analyzing changes in EMT markers and cytosolic/nuclear area, cells were seeded at a density of 0.04×10^6^/ml in 0.3 ml media in an 8 well glass-bottom chamber coated with Type 1 collagen (Purecol) (100 ug/mL). For assessing nuclear translocation of p65, cells were seeded at a density of 0.2×10^6^/ml in 0.3 ml media in a coated chamber as described above. The following day, the cells were treated as before for 72 hrs (for markers) or 30 min (for p65), fixed with 4% paraformaldehyde (Electron Microscopy Sciences) for 15 min at 37°C, washed in DPBS (Gibco) and permeabilized/blocked with blocking solution (0.3% TritonX-100 and 3% BSA containing DPBS) for 1 hr at room temperature (RT). Cells were then stained with primary antibodies against E-Cad, N-Cad, Fn, Vim, or p65 diluted in the blocking solution supplemented with 1% goat serum and incubated at 4°C for overnight. The dilutions used for the primary antibodies and their sources are presented in Table 2. The next day, cells were washed in DPBS and stained with AF488 or AF568 fluorochrome conjugated goat anti-mouse and anti-rabbit secondary antibodies (dilution 1:500, Invitrogen), as well as DAPI (D9542, Sigma Aldrich) and/or Phalloidin (P1951, Sigma Aldrich). Cells were imaged using a 63x or 20X objective lens on a Zeiss LSM880 Airyscan confocal microscope. Cells in three to five different fields across the well were captured randomly per condition in each experiment.

**Table 2.** Antibody information for western blotting and IF.

| **Antibody against** | **Species** | **Dilution**<br>**WB** | **Diluti**<br>**on IF** | **Distributor/Catalog**<br>**Number** |
| --- | --- | --- | --- | --- |
| Vim | Rabbit | 1:500 | 1:500 | Cell Signaling Technology,<br>5741S |
| E-Cad | Mouse | 1:10,000 | 1:500 | BD Biosciences, 610181 |
| N-Cad | Mouse | 1:500 |  | Santa Cruz Biotechnology,<br>sc-59987 |
| N-Cad | Rabbit |  | 1:500 | Proteintech 22018-1-AP |
| Snail | Rabbit | 1:500 | 1:500 | Cell Signaling, 3879S |
| Twist | Mouse | 1:650 | 1:500 | Santa Cruz Biotechnology,<br>sc- 81417 |
| Fibronectin | Mouse | 1:500 | 1:500 | Santa Cruz Biotechnology,<br>sc-271098 |
| NFkB p65 | Rabbit |  | 1:100 | Invitrogen, 14-6731-81 |
| P-SMAD3 | Rabbit | 1:1000 |  | Boster Bio P00059-1 |
| P-p38MAPK<br>(Thr180/Tyr182) | Rabbit | 1:1000 |  | Cell Signaling Technology,<br>4511 |
| P-ERK (Thr202/Tyr204) | Rabbit | 1:2000 |  | Cell Signaling Technology,<br>4370 |
| P-JNK (Thr183/Tyr185) | Rabbit | 1:1000 |  | Cell Signaling Technology,<br>9251 |
| P-AKT (Thr308) | Rabbit | 1:1000 |  | Cell Signaling Technology,<br>4056 |
| SMAD3 | Rabbit | 1:1000 |  | Cell Signaling Technology,<br>9513 |
| p38MAPK | Rabbit | 1:1000 |  | Cell Signaling Technology,<br>9212 |
| ERK | Rabbit | 1:1000 |  | Cell Signaling Technology,<br>4695 |
| JNK | Rabbit | 1:1000 |  | Cell Signaling Technology,<br>9252 |
| AKT | Rabbit | 1:1000 |  | Cell Signaling Technology,<br>4691 |
| GAPDH | Mouse | 1:1000 |  | Santa Cruz Biotechnology,<br>sc-166574 |
| Alpha-tubulin | Mouse | 1:2000 |  | Proteintech, HRP-66031 |
| GFP | Rabbit | 1:1000 |  | Abcam, ab290 |

### Western Blotting

Cells following treatments in a six well plate were lysed using radio immunoprecipitation assay (RIPA) buffer supplemented with Halt protease inhibitor (ThermoFisher #87786). Collected lysates were clarified by centrifugation at 13,000 rpm for 15 min at 4°C. The protein contents of each sample were then estimated using the BCA Protein Assay Kit (ThermoFisher #23225). An equal amount of protein was resolved by 10% SDS–PAGE transferred onto PVDF or nitrocellulose membranes (Millipore), blocked with 5% nonfat dry milk for 1 hr and probed with primary Abs against phospho- and total p44/42 MAPK (ERK1/2), p38MAPK, JNK, AKT, SMAD3, GAPDH, and alpha tubulin. The dilutions used for the primary antibodies and the sources are described in Table 2. Bands were visualized using HRP-conjugated secondary antibodies (dilution 1:8000, Jackson ImmunoResearch), SuperSignal™ West Pico PLUS Chemiluminescent Substrate, and a C600 digital imaging system (Azure). Integrated density for each band was measured using ImageJ software.

### Transwell Assay

To quantify the ability of CM to induce neutrophil migration, transwell migration chambers with membrane inserts of 3 μm diameter pore size (Greiner Bio-One) were used. Both inserts and bottom wells were coated with 2% tissue culture grade BSA for 1 hr at 37°C to prevent strong neutrophil adhesion. Coated inserts and wells were rinsed with DPBS twice to remove residual BSA. Freshly isolated neutrophils resuspended in Ca^++^, Mg^++^ HBSS (Gibco) at a density of 4×10^6^/ml were seeded onto the inserts (100 μl) and placed in a 24-well plate. Control chemoattractants or CM were gently added in 600 μl volume to the bottom wells of the 24-well plate. For the neutralization assay, CM or chemoattractant was incubated with 5 µg/ml antibodies for 30 min at 37°C with gentle rotation before they were added to the bottom wells. Migration was allowed to take place in 37°C/5% CO_2_ for 2 hrs. The percentage of neutrophils migrated to the bottom chamber was calculated from the cell counts obtained using a hemocytometer.

### Cancer Cell Migration and Invasion Assay

Endpoint migration and invasion assays were performed using a transwell system in 24-well plates as described before (6). Briefly, fluoroBlock™ filter inserts (351152, Corning) with 8 μm pore size were left uncoated for the migration assays or coated for the invasion assays with 0.2 mg/ml type I bovine collagen PureCol® of which the pH was neutralized to 7.2 – 7.4 using sodium bicarbonate. Cell lines were serum starved for 24 hrs and then seeded onto the inserts at 0.5×10^5^ cells/ml in 100 μl of serum free media. Cells were allowed to migrate toward the bottom chamber containing 500 μl of full-serum media as the chemoattractant or serum-free media as the negative control. After 24 hr incubation at 37°C/ 5% CO_2_, the inserts were transferred to a fresh 24-well plate with black walls containing 500 ml of 4 µM Calcein AM (Biotium) in Ca^++^, Mg^++^ HBSS per well. Cells were incubated for 1 hr at 37°C, and the fluorescence reading of migrated/invaded cells was measured from the bottom at wavelengths of 495/515 nm (Excitation/ Emission) by a SpectraMax M5 Multi-Mode Microplate Reader (Molecular Devices). Fluorescence reading of the negative control was subtracted from the reading for full-serum media condition.

### ELISA Assay

The presence of neutrophil recruiting chemokines; CXCL1, CXCL2, CXCL8, and activating factor; TGF-β1 in the CM was quantified by ELISA through the Rogel Cancer Center Immunology Core. Samples were tested with dilutions in quadruplicate. The protein concentration for each target analyte was quantified by comparing the colorimetric signal from the sample with the individual standard curve generated by known concentrations of each protein.

### Image quantification and data representation

CellProfiler (ver.4.2.5) was used to segment cells with “IdentifySecondaryObjects” module and nuclei with “IdentifyPrimaryObjects” module based on “Minimum Cross-Entropy or Otsu” algorithms for intensity thresholding. Cytosol was segmented by subtracting nuclei from cell with “IdentifyTertiaryOjects” module. The integrated intensities of EMT markers were quantified with “MeasureObjectIntensity” module using “Minimum Cross-Entropy or Otsu” algorithms for intensity thresholding. Cytosolic area as presented in Fig. 1Bi was quantified from the maximum intensity projections (MIPs) of the z-stacks of phalloidin-TRITC and DAPI stained cells as in Fig. 1Aiii,iv. Nuclear area as presented in Fig. 1Bii was quantified from representative *z*-stacks of DAPI-stained nuclei as in Fig. 1Av,vi. Ratio of cortex to cortex free cytosolic signal intensities for E-Cad were calculated from representative z-stacks of E-Cad/Vim/DAPI-stained cells as in Fig. 1Av,vi. Signals of Fn, Vim, and N-Cad were quantified from a representative *z*-stack of individual images as in Fig. 1Bv,vi,vii,viii or the MIP of the z-stacks of individual images as in Fig. 5Civ-ix. For measuring nuclear translocation of p65, ratio of nuclear to cytosolic signal intensities for p65 were calculated from the MIPs of the z-stacks of phalloidin-TRITC/p65/DAPI-stained cells as in Fig. 4B.

### Gene expression dataset analysis

Havard Medical School LINCS (Library of Integrated Network-based Cellular Signatures) dataset ID:20348 was downloaded through the LINCS DB database website (March 2023). Cell lines were then annotated by their respective tumor subtype. Correlations between all genes in the dataset were calculated using R v.4.2.2, cor function and p-values adjusted using FDR method.

DepMap 23Q2 mRNA expression data (1450 cell lines) was downloaded through the DepMap Portal (https://depmap.org/portal) and analyzed using R v.4.2.2. Cell types were then annotated by their lineage (‘OncotreeLineage’) to identify differential correlations between breast cancer cell lines vs the entire database. All correlations were calculated using R and p-values adjusted using FDR method. The top 100 correlating genes were uploaded to the PANTHER Classification System API (https://www.pantherdb.org/, (73), and were analyzed for statistical overrepresentation test (PMID 30804569), using Reactome pathways annotation (74). Pathways with FDR adj.P<0.05 were plotted for their fold-enrichment using GraphPad Prism.

### Statistical Analysis

GraphPad Prism software was used for data plotting and conducting statistical analysis by tests that are described in the respective figure legends along with the size of the samples. Tests used included two-tailed paired t test, unpaired t test, 1-way ANOVA with Dunnett’s multiple comparisons test or 2-way ANOVA with Sidak’s or Turkey multiple comparisons test.

## ACKNOWLEDGEMENTS

We are thankful to Dr. Jing Yang, UCSD for kindly providing pWZL-Blast-twist construct. We acknowledge Dr. Michael Holinstat and Amanda Prieur from the Platelet Physiology and Pharmacology Core for providing blood draws for this study and thank Peilin Shen for neutrophil isolation and technical assistance. We thank the Immunology core at the University of Michigan for expertise in processing ELISA samples. We acknowledge S. Arya (University of Michigan) for their intellectual contributions. We thank all the members of the Parent laboratory for their valuable suggestions.

## AUTHOR CONTRIBUTIONS

Conceptualization: SS, EC, PAC, CAP

Methodology: SS, EC, JS, KO

Investigation: SS, EC, JS, KO

Visualization: SS, EC, JS, KO

Funding acquisition: CAP

Supervision: PAC, CAP

Writing – original draft: SS, EC

Writing – review & editing: SS, EC, JS, KO, PAC, CAP

## SUPPLEMENTARY MATERIALS

**Figure S1.**
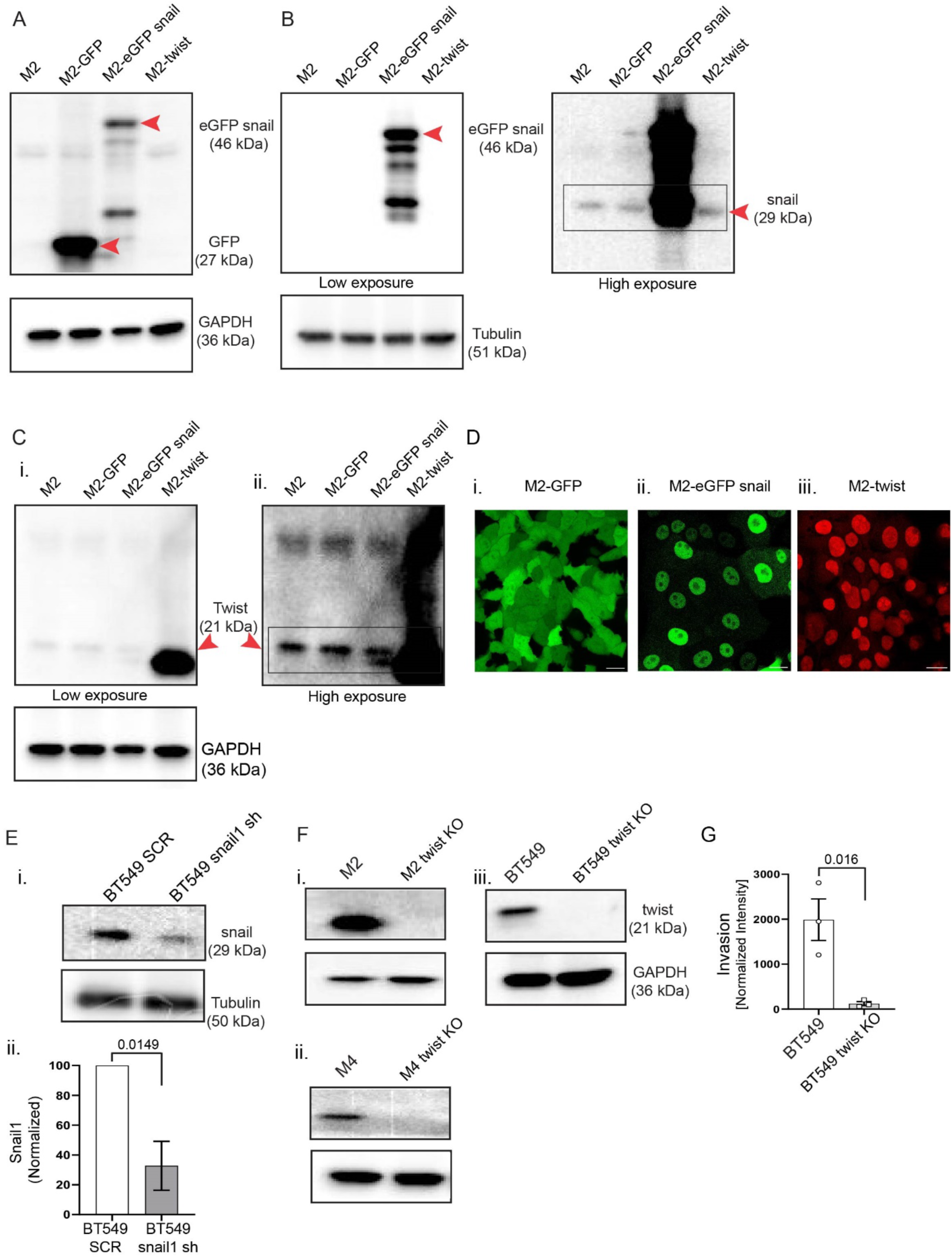
Snail and twist expression in engineered cell lines. **A-C.** Representative western blots showing the expression of GFP or eGFP-snail with anti-GFP (A) or anti-snail Ab (B; low (left) or high (right) exposures) and twist with anti-twist Ab (C; low (left) or high (right) exposures) in the respective cell lines (n=2). **D.** Representative IF images (n=2) showing MIP images for GFP in live M2 GFP (i), M2-eGFP snail cells (ii) or of fixed M2-twist cells stained for twist Ab (red) (iii). Scale bar 20 µm. **E.** Representative western blot showing the expression of snail in BT549 snail sh compared to SCR cell lines (i). Bars showing band intensities of snail normalized to the loading control (mean values +/− SEM from n=3). **F.** Representative western blot showing the expression of twist from M2 (i), M4 (ii), and BT549 (iii) twist KO cell lines (n=2). **G.** Graphs showing the invasion of BT549 WT and twist KO cell lines (mean values +/− SEM from n=3). P values were determined using unpaired t test (Eii and G).

**Figure S2.**
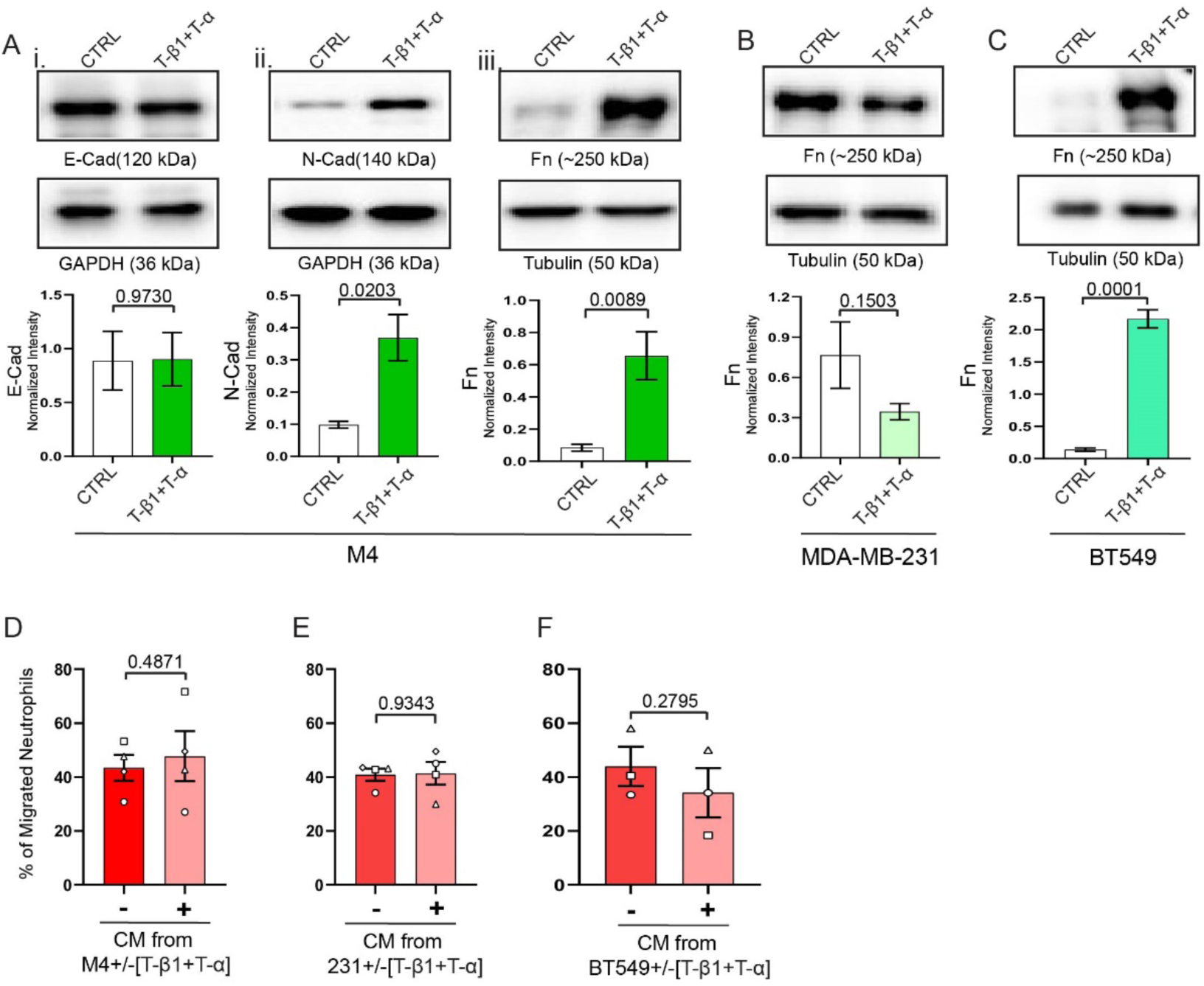
TGF-β1/TNF-α treatment does not boost neutrophil recruiting activity of TNBCs. **A-C.** Top: Representative western blots showing E-Cad (Ai), N-Cad (Aiii) or Fn (Aiii, B,C) expression of CTRL or T-β1/T-α treated M4, MDA-MB-231 and BT549 cells. Bottom: Graphs showing band intensities of the respective markers normalized to loading controls are presented (mean values +/− SEM from n=3-4). **D-F.** Graphs depicting the percentage of neutrophils that migrated into the bottom chamber of transwells containing equal volume of CM from CTRL or T-β1/T-α treated TNBCs (mean +/− SEM form n=3-4). Each dot represents response of neutrophils from an independent donor. P values were obtained using Unpaired (A,D) or Paired (B,C) t test.

**Figure S3.**
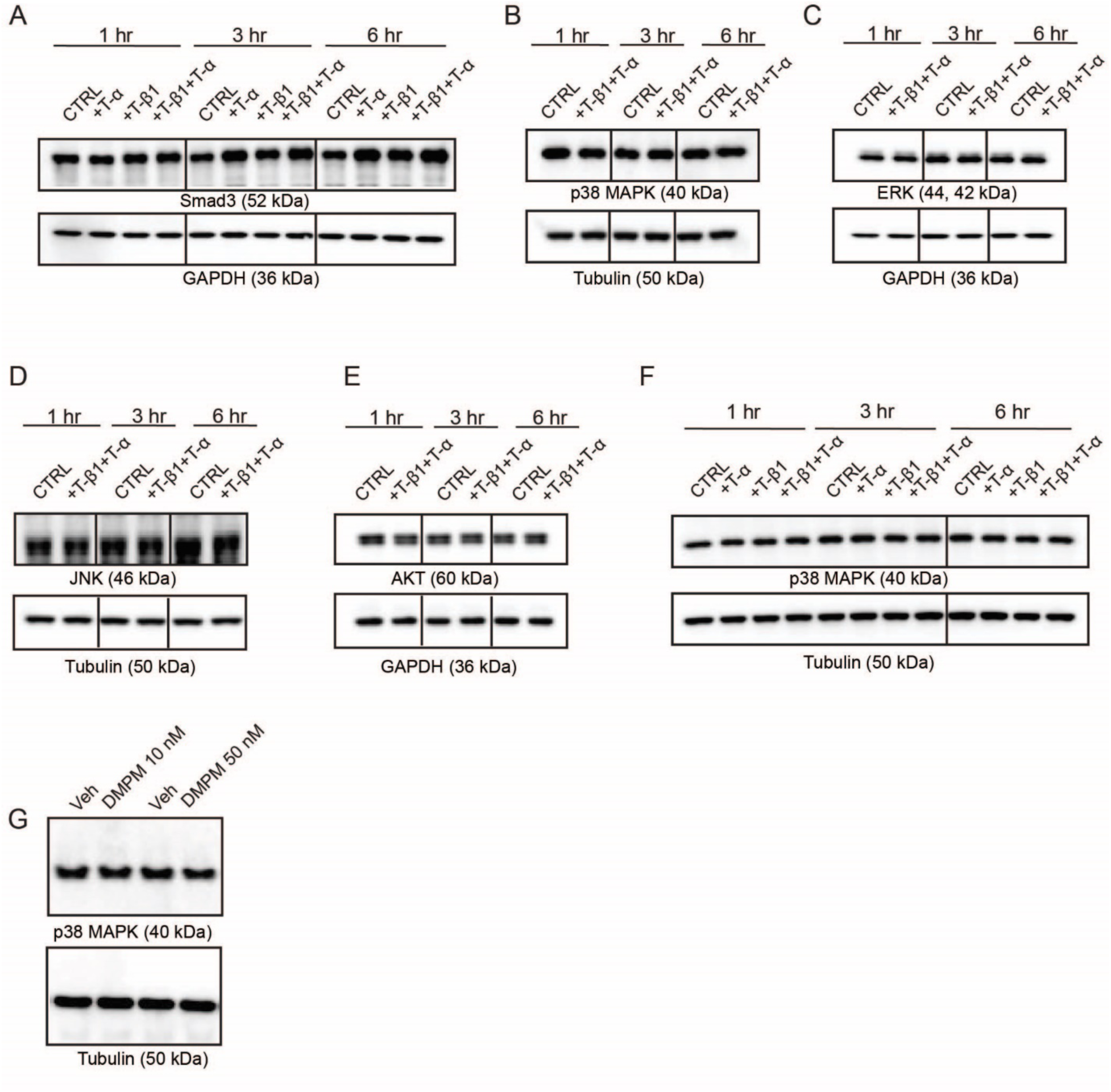
TGF-β1/TNF-α treatment does not affect total protein levels of the activated signaling pathways in M2 cells. **A-F.** Representative western blots showing Smad3 (A), p38MAPK (B, F), ERK (C), JNK (D) or AKT (E) expression of CTRL M2 cells or M2 cells treated with T-β1 or T-α or T-β1+T-α over the course of 1, 3, and 6 hrs of treatments (n=3). **G.** Representative western blots showing p38MAPK expression of M2 cells pre-treated with DMPM or vehicle control and stimulated with T-β1+T-α for 72 hrs (n=3).

**Figure S4.**
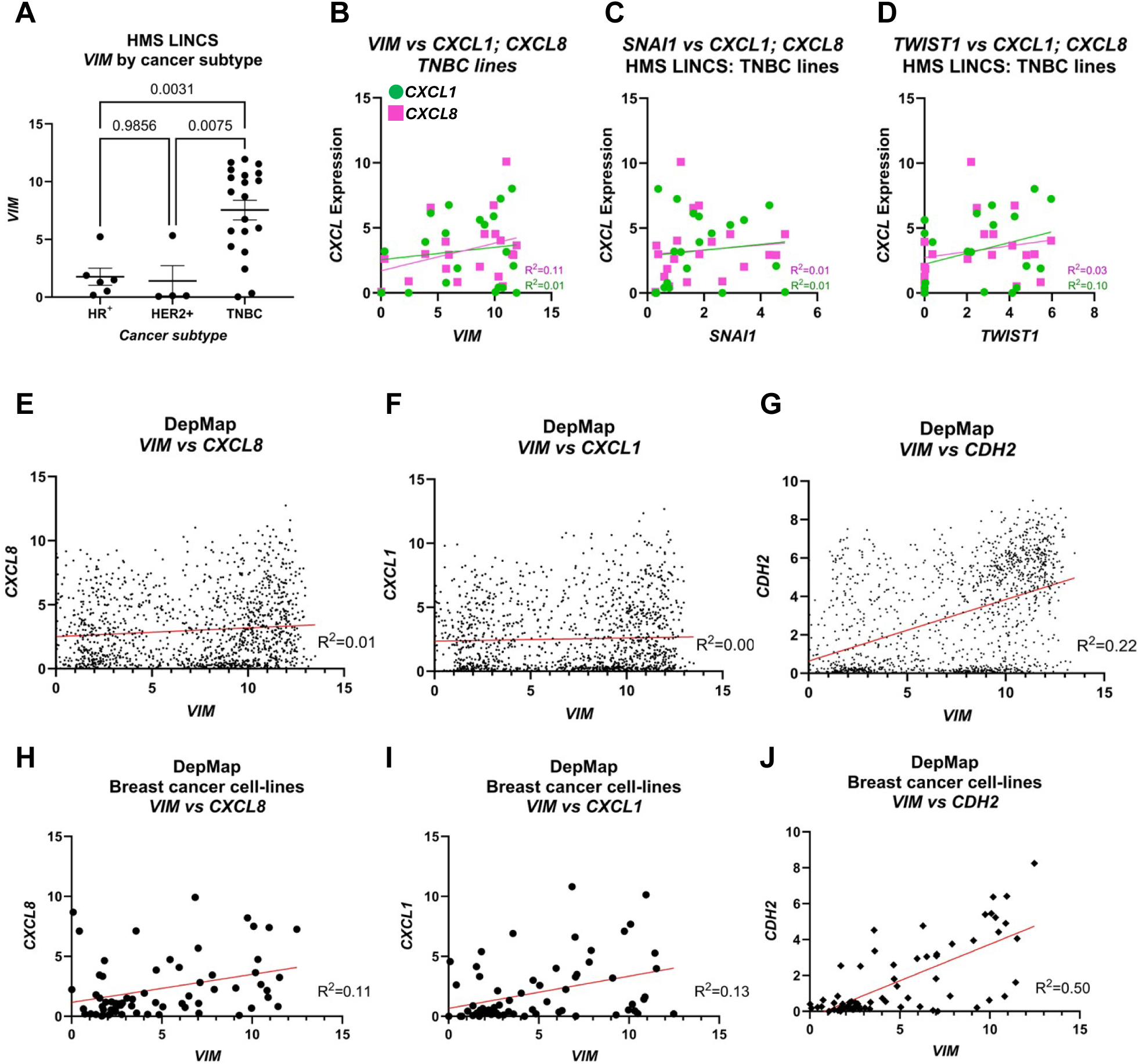
Analysis of gene expression datasets identifies no association between *CXCL8* or *CXCL1* and *VIM* expressions. **A.** Expression level of *VIM* in HR+, HER2+, or TNBC cell lines available in the LINCs dataset. **B-D.** Individual gene correlation between *CXCL8* or *CXCL1* against EMT markers *VIM* (B), *SNAI1* (C) and *TWIST1* (D) in 20 TNBC cell lines in the LINCs dataset. **E-G.** Correlation between *VIM* and *CXCL8* (E), *CXCL1* (F) or *CDH2* (G) across all available cell lines in the DepMap dataset. **H-J.** Correlation between *VIM* and *CXCL8* (H), *CXCL1* (I) or *CDH2* (J) across breast cancer cell lines in the DepMap dataset.

**Table S1.** TGF-β1/TNF-α treatment does not induce TGF-β1 secretion from M2 cells. Table showing the amount (pg/ml) of TGF-β1 secreted by CTRL or T-β1/T-α treated M2 cells from three independent experiments.

| Treatment condition | T-β1 in CM (pg/ml) |
| --- | --- |
| CTRL | 0 |
| T-β1+T-α | 0 |
| CTRL | 0 |
| T-β1+T-α | 0 |
| CTRL | 4 |
| T-β1+T-α | 5 |

Other supplementary file

